# Loss of the DYRK1A protein kinase results in reduction of ribosomal protein genes expression, ribosome mass and translation defects

**DOI:** 10.1101/2021.01.21.427583

**Authors:** Chiara Di Vona, Laura Barba, Roberto Ferrari, Susana de la Luna

## Abstract

Ribosomal proteins (RPs) are evolutionary conserved proteins that are essential for protein translation. RP expression must be tightly regulated to ensure the appropriate assembly of ribosomes and to respond to the growth demands of cells. The elements regulating the transcription of RP genes (RPGs) have been characterized in yeast and *Drosophila*, yet how cells regulate the production of RPs in mammals is less well understood. Here, we show that a subset of RPG promoters is characterized by the presence of the palindromic TCTCGCGAGA motif and marked by the recruitment of the protein kinase DYRK1A. The presence of DYRK1A at these promoters is associated with enhanced binding of the TATA-binding protein, TBP, and it is negatively correlated with the binding of the GABP transcription factor, establishing at least two clusters of RPGs that could be coordinately regulated. However, DYRK1A silencing leads to a global reduction of RPGs, mRNA pointing at DYRK1A activities beyond those dependent on its chromatin association. Significantly, cells in which DYRK1A is depleted have reduced RP levels, fewer ribosomes, reduced global protein synthesis and smaller size. We therefore propose a novel role for DYRK1A in coordinating the expression of genes encoding RPs, thereby controlling cell growth in mammals.

## INTRODUCTION

Ribosomes are cellular machines that translate mRNA into protein, and in mammals they are formed by the large 60S subunit and the small 40S subunit. The 60S subunit is comprised of the 5S, 5.8S and 28S rRNAs associated with 52 ribosomal proteins (RPs), and the smaller 40S subunit is made up of the 18S rRNA plus 35 RPs. Ribosome biogenesis is a complex process that involves more than 200 different factors: rRNAs, small nucleolar RNAs, and canonical and auxiliary RPs (1). The three RNA polymerases (Pol) participate in the transcription of the ribosomal components, with Pol I responsible for transcribing the 28S, 18S and 5.8S rRNAs, Pol III transcribing the 5S rRNAs and Pol II responsible for the transcription of all the protein coding genes involved in ribosome biogenesis, including the RP genes (RPGs). Therefore, the coordinated expression of these components is required to ensure the correct assembly and proper functioning of ribosomes. Indeed, dysregulation of ribosome biogenesis is associated with a group of human diseases that are collectively known as ribosomopathies (2) and moreover, alterations to RP expression contribute to cancer cell growth (3).

The coding sequences of RPGs have been highly conserved over evolution, unlike the features of their promoters and the machinery involved in their transcriptional regulation. As such, RPGs are organized into operons in prokaryotes (4), whereas the situation is much more complex in the case of eukaryotes, with multiple genes widely scattered across the genome (5). The main elements involved in the transcriptional regulation of RPGs have been characterized thoroughly in *Saccharomyces cerevisiae* (6), in which the Repressor Activator Protein 1 (Rap1p) and the Fhl1p forkhead transcription factor (TF) are constitutively bound to the RPG promoters, coordinating RPG expression (7). In higher eukaryotes, most studies have focused on the differential enrichment of TF binding motifs within RPG promoters (8–11). In particular, several DNA sequences are found over-represented in human RPG promoters. The polypyrimidine TCT motif is found close to the transcription start site (TSS) of RPGs, and it is thought to play a dual role in the initiation of both transcription and translation (12). This motif is recognized by the TATA-box Binding Protein (TBP)-Related Factor 2 (TRF2) in *Drosophila* (13, 14), yet it remains unclear whether there is functional conservation with its human TBP like 1 (TBPL1) homolog. Around 35% of the RPG promoters contain a TATA-box in the -25 region and an additional 25% contain A/T-rich sequences in this region (15). Other motifs frequently found are those for SP1, the GA-Binding Protein (GABP) and the Yin Yang 1 (YY1) TFs (15). In addition, the E-box TF MYC is a key regulator of ribosomal biogenesis, enhancing the expression of RPGs (16). Finally, a *de novo* motif (M4 motif) was found enriched in human and mouse RPG promoters (8). RPG mRNA expression displays tissue- and development-specific patterns, both in human and mouse (5,17,18). Hence, RPG expression could be regulated by specific combinations of TFs in different organisms and/or physiological conditions.

The M4 motif matches the palindromic sequence that is bound by the dual-specificity tyrosine-regulated kinase 1A (DYRK1A) protein kinase (19). DYRK1A fulfills many diverse functions by phosphorylating a wide range of substrates (20, 21), and it is a kinase with exquisite gene dosage dependency. On the one hand, DYRK1A overexpression in individuals with trisomy 21 has been associated to several of the pathological symptoms associated with Down syndrome (DS) (22). On the other hand, *de novo* mutations in one *DYRK1A* allele cause a rare clinical syndrome known as DYRK1A haploinsufficiency syndrome (OMIM#614104) (23, 24) and references therein). DYRK1A has also been proposed as a pharmacological target for neurodegenerative disorders, diabetes and cancer (21,25,26). We have shown that DYRK1A is a transcriptional activator when recruited to proximal promoter regions of a subset of genes that are enriched for the palindromic motif TCTCGCGAGA (19). DYRK1A phosphorylates serine residues 2, 5 and 7 within the C-terminal domain (CTD) of the catalytic subunit of Pol II (19). This activity takes over that of the general TF P-TEFb at gene loci involved in myogenic differentiation (27). The interaction of DYRK1A with the CTD depends on a run of histidine residues in its non-catalytic C-terminus, which also promotes the nucleation of a phase-separated compartment that is functionally associated with transcriptional elongation (28). Here, we have analyzed the occupancy of RPG promoters by DYRK1A in more depth, performing a comprehensive analysis of the promoter occupancy by other factors whose binding motifs are differentially enriched in human RPG promoter regions. Our results indicate that most of these factors are found at almost all RPG promoters, irrespective of the presence of their cognate binding sites. By contrast, DYRK1A associates with a subset of human and mouse RPG promoters that contain the TCTCGCGAGA motif. Moreover, physiological levels of DYRK1A are required to maintain RPG transcript levels independently of the binding of DYRK1A to their promoters, and this effect could at least in part contribute to the global reduction in ribosome mass and protein synthesis when DYRK1A is silenced. Therefore, our results expand the functional spectrum of the DYRK1A kinase, indicating that it contributes to the regulation of cell growth in mammalian cells.

## RESULTS

### DYRK1A is recruited to the proximal promoter regions of canonical RPGs

Our prior analysis of DYRK1A recruitment to chromatin showed an enrichment in gene ontology terms related to ribosome biogenesis and translational regulation (19). In addition, a *de novo* motif analysis found a sequence similar to the DYRK1A-associated motif to be over-represented in the promoter regions of human RPGs (8). Accordingly, we examined whether DYRK1A was recruited to human RPGs, considering the genes encoding the 80 canonical RPs plus 10 paralogues (5); a new naming system for RPs has been proposed (29) and the equivalences to this nomenclature are listed in Table Supplementary 1. In experiments on different human cell lines, including glioblastoma T98G, bone osteosarcoma U2OS and cervical adenocarcinoma HeLa cells (Figure 1A), genome-wide ChIP followed by deep sequencing (ChIP-Seq) showed DYRK1A bound upstream of the TSS and in general, within 500 bp of the TSS of a subset of RPGs (Figure 1A). No ChIP signal was detected at either the gene bodies or at the transcriptional termination sites (TTS; Figure Supplementary 1). To validate the recruitment of DYRK1A to RPG promoters, genome-wide DYRK1A occupancy was assessed in both control cells and in cells where the levels of DYRK1A were depleted by lentiviral delivery of a shRNA targeting DYRK1A. A clear reduction in chromatin-associated DYRK1A was observed at target RPG promoters, reflecting the specific depletion of this kinase (Figure 1B; Figure Supplementary 1). Finally, the presence of DYRK1A at specific promoter regions of RPGs was further confirmed in independent ChIP-qPCR experiments (Figure 1C).

**Figure 1.**
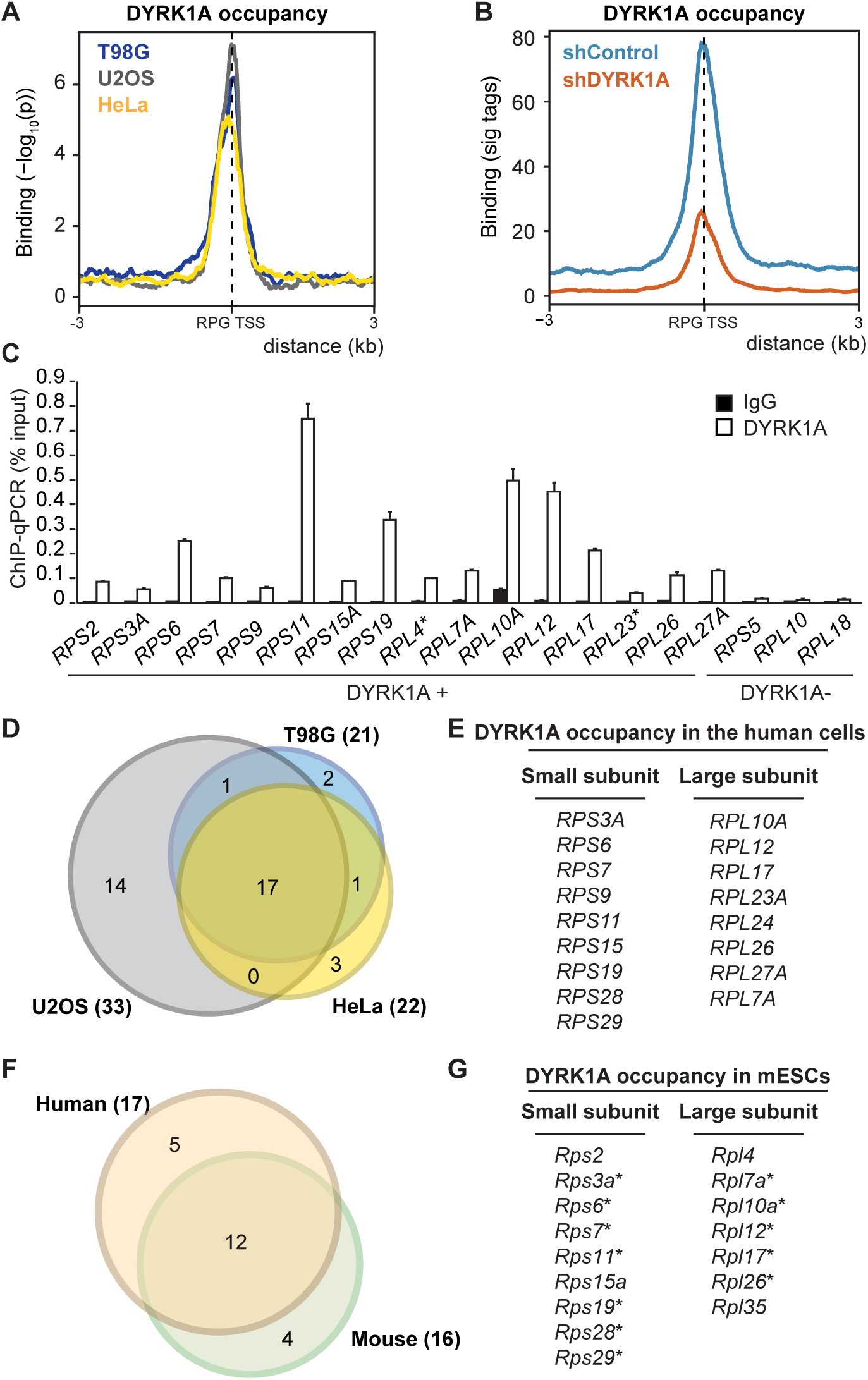
DYRK1A occupies a subset of RPG promoters. (**A**) Density plot showing the distribution of DYRK1A in T98G, U2OS and HeLa cells relative to the TSS of all human RPGs. The *y*-axis represents the relative protein recruitment (-log_10_ Poisson *p*-value) and the offset was set to ± 3 kb from the TSS. (**B**) Density plot showing DYRK1A occupancy relative to the TSS of all human RPGs, comparing shControl and shDYRK1A T98G cells (blue and orange lines, respectively). The *y*-axis represents the relative protein recruitment quantified as significant (sig) ChIP-Seq tags. The offset was set to ± 3 kb from the TSS (see Figure Supplementary 1B for representative examples). (**C**) Validation of selected targets by ChIP-qPCR, representing the data as a percentage of the input recovery (mean ± SD of three technical replicates). See Key Resources Table for primer sequences. (**D**) Venn diagram showing the overlap of DYRK1A-associated RPG promoters in T98G, U2OS and HeLa cells. (**E**) List of DYRK1A-positive RPGs common to the three human cell lines. (**F**) Venn diagram indicating the overlap between common DYRK1A-bound RPGs in the human cell lines and mESCs. (**G**) List of RPGs with DYRK1A detected at their promoters in mESCs. The asterisk indicates the coincident occupancy in the three human cell lines analyzed.

Around 25% of the promoters of all RPGs were occupied by DYRK1A in the different human cell lines analyzed, with considerable overlap among them (Figure 1D and E). No association with any particular ribosomal subunit was observed, since DYRK1A occupancy was detected similarly at the promoters of RPGs from both the large and small ribosome subunits (Figure 1E). Furthermore, DYRK1A ChIP-Seq data from mouse embryonic stem cells (mESCs) also showed the presence of this kinase at the promoters of a subset of mouse RPGs, which coincided well with the DYRK1A-positive subset in human cells (Figure 1F and G). Together, these results indicate that DYRK1A is recruited to proximal promoter regions of a subset of RPGs in different human and mouse cell lines, suggesting that the association of DYRK1A with these RPGs at the chromatin level represents a general and conserved function for the kinase.

### The TCTCGCGAGA motif marks the subset of RPG promoters positive for DYRK1A

Around 25% of the human RPG promoters contain a TCTCGCGAGA motif (Figure 2A and B), and DYRK1A bound to these in one, two or in all the human cell lines analyzed, except for *RPL10* and *RPS5* and the RPG paralogs *RPL10L*, *RPL22L1* and *RPS4Y2* (Table Supplementary 1), which are not expressed in any of these cell lines. Likewise, DYRK1A was almost exclusively detected at RPG promoters containing the TCTCGCGAGA motif in mESCs (Table Supplementary 2), as further evidence of its functional conservation. Therefore, the TCTCGCGAGA motif appears to be a distinctive feature of DYRK1A-binding RPG promoters in different cell lines.

**Figure 2.**
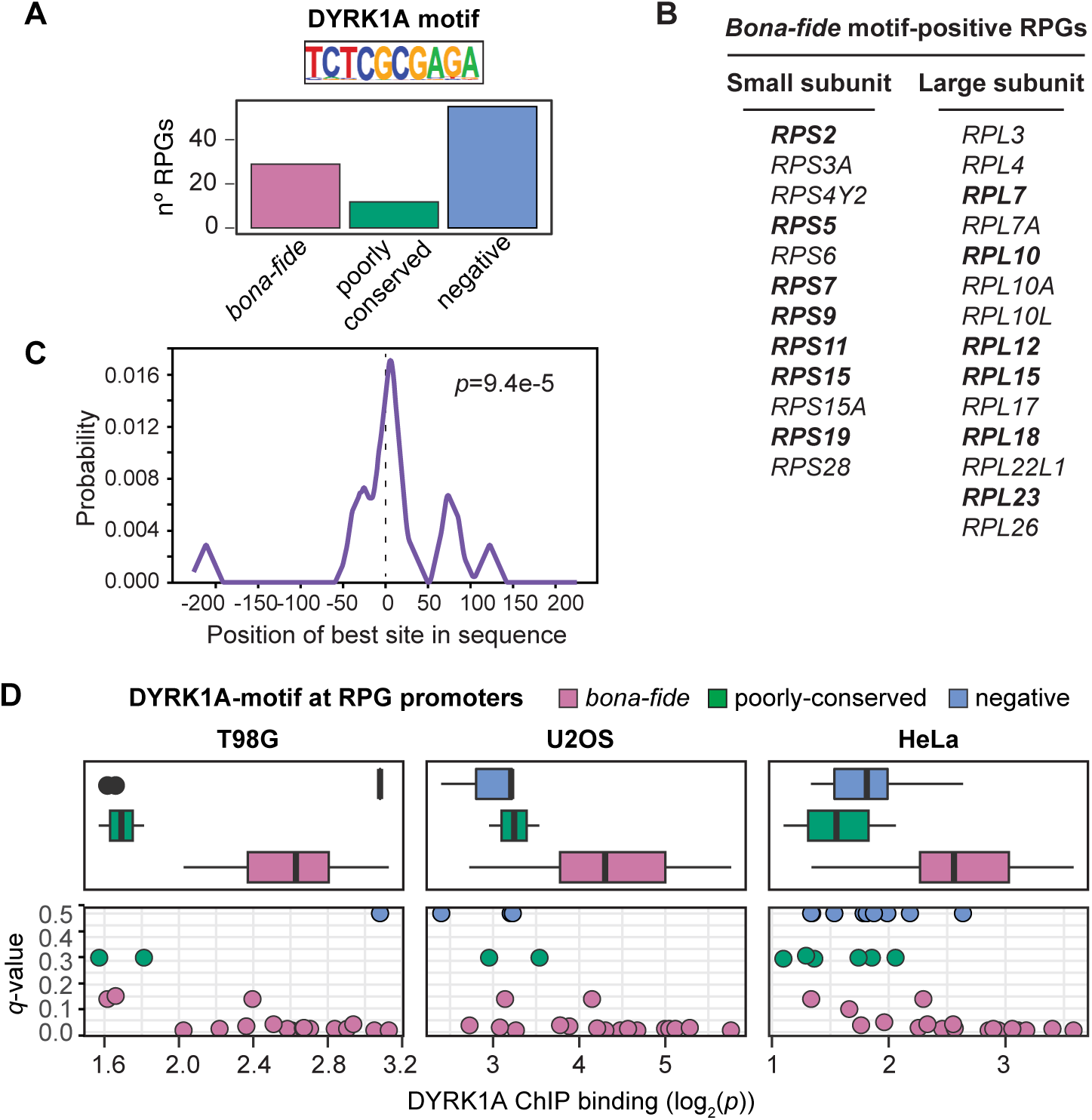
The TCTCGCGAGA motif correlates with significant DYRK1A binding at human RPG promoters. (**A**) Distribution of the TCTCGCGAGA motif (DYRK1A motif) in human RPGs (*bona-fide*: *p*-value < 10^-4^; poorly conserved: 10^-4^ < *p*-value < 3×10^-4^). The DYRK1A motif associated logo is included. (**B**) List of human RPGs with the *bona-fide* palindromic motif detected in their promoters. The RPG promoters containing more than one DYRK1A motif are highlighted in bold. (**C**) CentriMo plot showing the distribution of the DYRK1A motif for the RPG promoters that bind DYRK1A. The solid curve shows the positional distribution (averaged over bins of 10 bp-width) of the best site of the DYRK1A motif at each position in the RPG-ChIP-Seq peaks (500 bp). The *p*-value for the central enrichment of the motif is included. (**D**) Box plot and scatter plots (dots represent individual RPGs) showing the correlation between DYRK1A ChIP binding (log2 *p*-value, *x*-axis) and the conservation of the DYRK1A motif (*q*-value, *y*-axis) in T98G, U2OS and HeLa cells. The colors represent the motif classification indicated.

A central model motif analysis (CMEA, 30) showed the TCTCGCGAGA motif to be positioned precisely around the center of the DYRK1A-associated peaks within the RPG promoters (Figure 2C), suggesting it serves as a platform for DYRK1A recruitment. The kinase was also found to associate with a small number of RPG promoters without this palindromic sequence (Table Supplementary 1), although its detection was poorer in these cases than when the RPG promoter contained the TCTCGCGAGA motif (Figure 2D). Hence, this palindromic sequence appears to be a major determinant for DYRK1A efficient recruitment to RPGs. Given that the variability in DYRK1A occupancy in the different cell lines was mostly restricted to the RPG promoters without the TCTCGCGAGA motif, it is possible that DYRK1A associates with promoters containing this motif independently of the cell-type, whereas occupancy of other RPG promoters might be context-specific.

The TCTCGCGAGA motif has been shown to work as a promoter element that drives bidirectional transcription (31). In around 30% of RPGs, the TSS lies within a 500 bp window from the TSS of genes transcribed in the opposite direction and in these cases, the RPG is always transcribed more strongly (Supplementary File 1). However, no specific bias for the presence of the TCTCGCGAGA motif could be observed, since some RPGs with bidirectional transcription contained this motif (*RPL12*, *RPL15*, *RPL23A*) while others did not (*RPL34*, *RPL9*, *RPS18*; Supplementary File 1). The transcriptional repressor Zinc finger and BTB domain containing 33 (ZBTB33, also known as KAISO) binds directly to the TCTCGCGAGA motif *in vitro* when methylated (32). Indeed, this palindromic motif is included in the Jaspar database of curated TF binding profiles as a ZBTB33-motif (http://jaspar.genereg.net/matrix/MA0527.1/). ChIP-Seq experiments in T98G cells did detect ZBTB33 at the majority of RPG promoters containing the TCTCGCGAGA motif (Figure Supplementary 2A), with signals overlapping those of DYRK1A (Figure Supplementary 2B). Moreover, the TCTCGCGAGA motif is positioned centrally within the ZBTB33 bound regions of these RPGs (Figure Supplementary 2C). Hence, this promoter motif might not only favor DYRK1A recruitment but also, its interaction with other proteins.

### Low functional conservation of RPGs core promoter elements between *Drosophila* and humans

Next, we wondered whether the RPG promoters that bind DYRK1A were characterized by any other feature. Most studies on the regulation of RPG expression in higher eukaryotes have used *Drosophila* as a model system, identifying several sequence elements and TFs that regulate RPG transcription (12–14). One of them is the TCT motif that is a specific promoter element for the expression of RPGs (12). Using high confidence human TSS data, we scanned human RPG promoters and we found the TCT consensus sequence (YC+1TYTYY; Figure 3A) in 77 of the 86 promoters analyzed (Supplementary Table 3). However, we did not find any correlation between the TCT motif and the presence of DYRK1A at these promoters (Figure 3B). In *Drosophila*, the TCT element drives the recruitment of Pol II and consequently, the transcription of RPGs through the coordinated action of TRF2 but not of TBP, and the TFs Motif 1 Binding Protein (M1BP) and DNA Replication-related Element (DRE) Factor (DREF) (13, 14). Thus, we asked whether the transcriptional regulators of RPGs were functionally conserved between *Drosophila* and humans, and if so, how they might be related to the promoter association of DYRK1A. We analyzed the presence of the homologs of the fly TFs at human RPG promoters: TBPL1, the homolog of TRF2; Zinc Finger Protein 281 (ZNF281), the homolog of M1BP; and Zinc finger BED-type containing 1 (ZBED1), the homolog of DREF. This analysis showed that TBPL1 binds to the promoter of almost all RPGs (Figure 3C and D; Supplementary Table 3), similar to its behavior in *Drosophila* (13). ZNF281 was only detected at 7 RPG promoters (Figure 3C; Supplementary Table 3), although the sequence motif enriched within the ZNF281 ChIP-Seq dataset differed from the M1BP binding motif (http://jaspar.genereg.net/matrix/MA1459.1/; 9) (Figure 3E) and thus, we cannot assume that M1BP and ZNF281 are fully functional homologs. ZBED1 binds to several human RPG promoters and gene bodies (Figure 3C; Supplementary Table 3), and the motif enriched in the ChIP-Seq dataset partially overlaps with the *Drosophila* DREF motif (http://jaspar.genereg.net/matrix/MA1456.1/; Figure 3E). However, no enrichment for this motif was found within the human RPG promoters. ZBED1 may recognize the TCTCGCGAGA motif (33), yet the CMEA analysis did not find a unimodal and centered distribution of the TCTCGCGAGA motif within the ZBED1 ChIP-peaks (Figure Supplementary 3A). Hence, ZBED1 did not appear to bind directly to the TCTCGCGAGA motif, consistent with data suggesting that the TCTCGCGAGA motif and the human DRE are distinct regulatory elements (31). Finally, ZBED1 binding to human RPGs showed no particular correlation with the presence of DYRK1A (Figure Supplementary 3B). Together, these data suggest there is little functional conservation of the core promoter elements of RPGs between *Drosophila* and humans, either in *cis* or *trans*. Furthermore, the association to chromatin of the TFs homologs to those identified in *Drosophila* does not appear to depend on the presence of the TCTCGCGAGA motif or the binding of DYRK1A to human RPG promoters.

**Figure 3.**
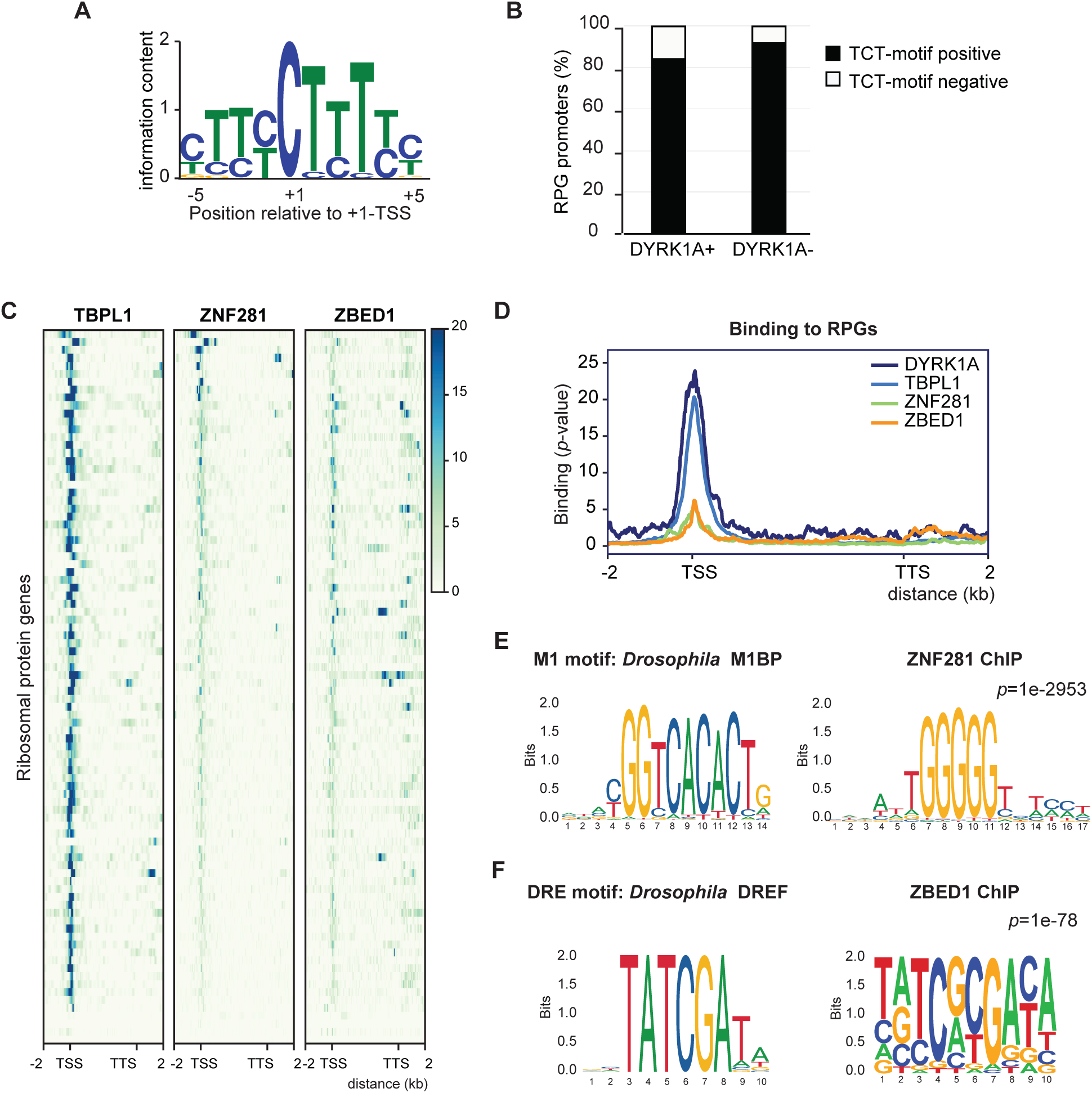
Analysis of *Drosophila* RPG expression regulators in mammalian cells. (**A**) Sequence logo of the human TCT motif found in human RPGs (see Materials and Methods). (**B**) Percentage of TCT positive or negative human RPGs promoters distributed according to the presence of DYRK1A. (**C**) The heatmaps show TBPL1 (K562, ENCODE dataset ENCSR783EPA), ZNF281 (HepG2, ENCODE dataset ENCSR403MJY) and ZBED1 (K562, ENCODE dataset ENCSR286PCG) occupancy of all human RPGs. The genomic region considered is shown on the *x*-axis. (**D**) Density plot of the occupancy of the transcription factors shown in the panel **C** together with DYRK1A. (**E**) Sequence of the *Drosophila* M1 motif (Jaspar: http://jaspar.genereg.net/matrix/MA1459.1/) and of the DNA motif enriched in ZNF281-bound regions. (**D**) Sequence of the DRE motif in *Drosophila* (Jaspar: http://jaspar.genereg.net/matrix/MA1456.1/) and of the motif enriched in ZBED1-bound regions.

### GABP and DYRK1A are differentially distributed at RPG promoters

We then asked whether DYRK1A recruitment was associated with the presence of TFs whose binding sites are known to be overrepresented at human RPG promoters, such as TBP, MYC, SP1, GABP and YY1. However, instead of using motif occurrence as in previous studies that analyzed RPG promoter architecture (10, 15), we took advantage of genome-wide mapping data for each of the TFs. Thus, though TBP was predicted to be differentially enriched at human RPG promoters based on the presence of the TATA-box (15); Supplementary Table 3), the analysis of TBP occupancy found TBP bound to nearly all human RPG promoters (93%) in the different cell lines (Figure 4A and B; Supplementary Table 3). Hence, TBP appears to be a general component of the human RPG transcriptional machinery, irrespective of the presence of a TATA consensus, a TATA-like sequence or the complete absence of such motif (Figure 4C). Consistent with a role for TBP in the assembly of the pre-initiation complex (PIC) at RPG promoters, the presence of the TFIID subunit TBP Associated Factor 1 (TAF1) was strongly correlated with that of TBP (Figure 4D; Supplementary Table 3). Notably, we observed stronger TBP binding at DYRK1A-enriched RPG promoters than at RPG promoters devoid of DYRK1A (Figure 4D and E), suggesting that these two factors might cooperate. The presence of YY1, SP1 and MYC was also detected in almost all RPG promoters irrespective of the presence of cognate binding sites (Figure Supplementary 4A; Supplementary Table 3), and an unbiased clustering analysis showed no differential distribution of any of these factors based on the presence of DYRK1A (Figure Supplementary 4B). By contrast, DYRK1A and GABP were distributed into one cluster that included those RPG promoters with high DYRK1A occupancy and low GABP presence (Figure 4F, cluster 1), and another in which promoters were depleted of DYRK1A but there was a strong GABP occupation (Figure 4F, cluster 2). Indeed, while the DYRK1A- associated TCTCGCGAGA motif was mostly overrepresented in cluster 1, the GABP-motif is a hallmark of cluster 2, and both motifs are distributed equally within the third cluster (Figure 4G). These results suggest that the presence of DYRK1A labels a specific subset of RPGs that might respond distinctly to those labeled by GABP.

**Figure 4.**
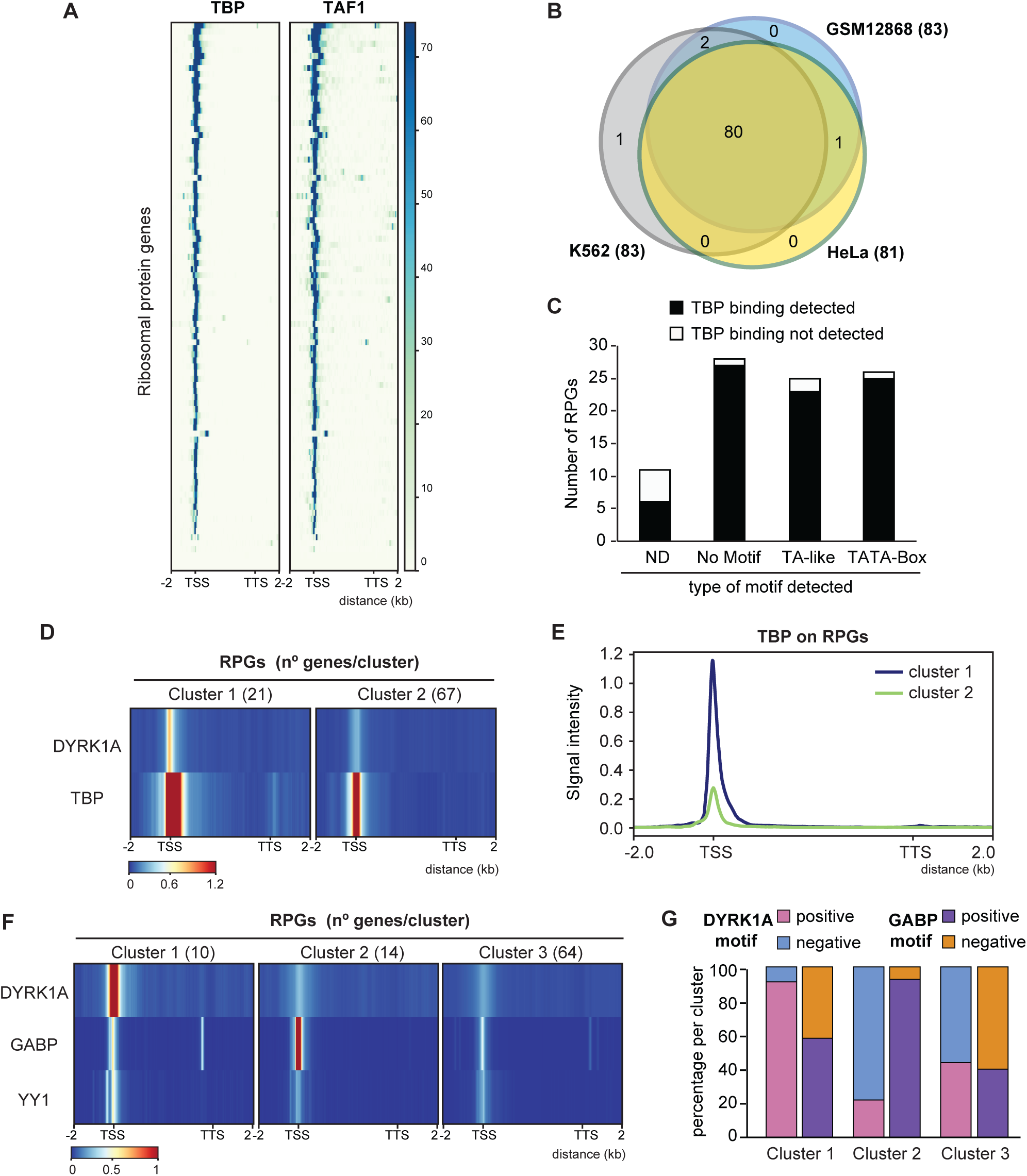
Differential recruitment of DYRK1A and GABP to RPG promoters. (**A**) The heatmaps show TBP and TAF1 (K562 cells, ENCODE datasets GSE31477 and ENCSR000BKS) occupancy in all human RPGs, showing the genomic region considered on the *x*-axis. (**B**) Venn diagram indicating the overlap of TBP occupancy at RPGs in different human cell lines (ENCODE dataset GSE31477). (**C**) Relationship of the presence of a TATA-box or TA-like sequences in human RPG promoters according to Perry et al (2005) and TBP binding in HeLa cells (ND, not determined). (**D**) Unbiased *k*-mean clustering of the average binding of DYRK1A (HeLa, this work) and TBP (HeLa, ENCODE dataset GSE31477) on all human RPGs. The color scale bar indicates the binding score and the genomic region considered is shown on the *x*-axis. (**E**) Metagene plot showing TBP occupancy relative to all human RPGs in HeLa cells according to the clusters shown in Figure 4D. The *y*-axis represents relative protein recruitment quantified as ChIP-Seq reads. (**F**) Unbiased *k*-mean clustering of the average binding of DYRK1A (T98G, this work), GABP and YY1 (SK-N-SH, ENCODE dataset GSE32465) to all RPGs. The color scale bar indicates the binding score and the genomic region considered is shown on the *x*-axis. (**G**) The histogram represents the percentage of RPG promoters containing a DYRK1A or a GABP motif in each of the clusters shown in Figure 4F.

### The expression of RPGs is sensitive to DYRK1A depletion

Based on the ability of DYRK1A to regulate transcription when recruited to chromatin (19), we wondered whether the transcription of RPGs might be modulated by DYRK1A. Indeed, the TCTCGCGAGA motif is a *cis*-element for the regulation of the expression of several RPGs, such as *RPL7A* (34), *RPS6*, *RPL10A*, *RPL12* (33) and *RPS11* (19). Furthermore, we demonstrated that the *RPS11* promoter responds to DYRK1A in a kinase- and motif-dependent manner (19). Globally, the expression of genes that have promoters occupied by DYRK1A is stronger than that of the genes in which DYRK1A is absent (Figure 5A). As described in *Drosophila* (13), the transcripts of most human RPGs were in the group of the top 5% most strongly transcribed genes in the cell lines analyzed (Figure Supplementary 5). The same tendency towards stronger expression of DYRK1A bound targets was observed when assessing RPGs, although the differences were not statistically significant in any of the cell lines analyzed (Figure 5B; Figure Supplementary 6). ChIP-Seq data of Pol II for RPGs revealed a profile corresponding to actively transcribed genes, with Pol II occupancy detected all along gene bodies, with the exception of those RPGs not expressed in the cell line analyzed (Figure 5C), and as an indication of transcriptionally engaged Pol II. No differences in the distribution of Pol II were found between the DYRK1A positive and negative RPGs (Figure 5C).

**Figure 5.**
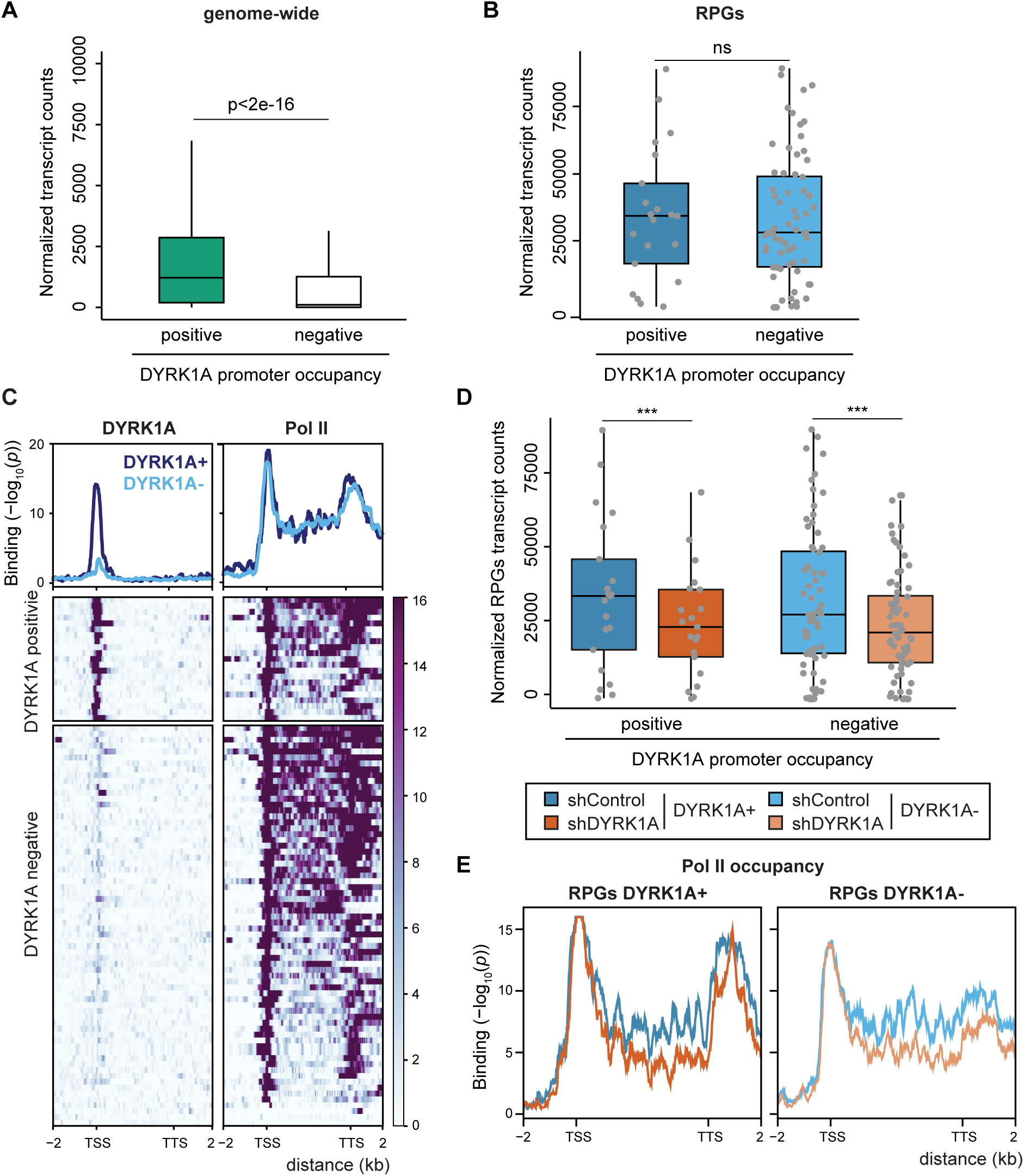
Depletion of DYRK1A causes a general reduction in RPG expression. (**A**) Box plot indicating the RNA levels of all human genes expressed in T98G cells (normalized counts) separated into two clusters depending on the presence of DYRK1A at their promoters (unpaired, two-tailed Student’s test). (**B**) Box plot indicating the RNA levels of all the human RPGs (normalized counts) separated into two clusters, as in Figure 5A (unpaired, two-tailed, Mann-Whitney test, ns=not significant). (**C**) *Bottom panel*, heatmap showing DYRK1A and Pol II chromatin occupancy relative to human RPGs depicted as metagenes, and separated into two clusters according to the presence/absence of DYRK1A at the promoters. The binding score (-log_10_ Poisson *p*-value) is indicated by the color scale bar. *Top panel*, density plot corresponding to the mean value of the heatmap (blue and light blue line correspond to DYRK1A positive or DYRK1A negative RPGs, respectively). The binding score (-log_10_ Poisson *p*-value) is indicated on the *y*-axis. (**D**) Box plot indicating the expression of RPGs (normalized counts) in T98G cells classified according to the presence (positive) or absence (negative) of DYRK1A at their promoters, and comparing cells infected with a lentivirus expressing a shRNA Control (blue and light blue, respectively) or shDYRK1A (orange and light orange, respectively; Wilcoxon matched-pairs signed ranks test, *** *p*<10^-5^). (**E**) Density plots corresponding to the mean value of Pol II binding in RPGs clusterized as positive and negative for DYRK1A binding in shControl (blue and light blue lines) and shDYRK1A conditions (orange and light orange lines). The binding score (-log_10_ Poisson *p*-value) is indicated on the *y*-axis.

We next assessed whether the absence of DYRK1A affected the expression of its target genes by comparing the RNA expression of cells infected with a control lentivirus or a lentivirus expressing a short hairpin (sh)RNA to DYRK1A, and we used *Drosophila* RNA for spike-in normalization. The majority of differentially expressed genes were downregulated in response to DYRK1A depletion (Figure Supplementary 7A). Indeed, these downregulated genes were enriched in the subset of genes with DYRK1A at their promoters (Figure Supplementary 7B), supporting a role for DYRK1A in transcriptional activation. A general reduction in RPG transcripts was observed in DYRK1A-silenced cells (Figure 5D), and the analysis indicated that RPGs whose promoters were occupied by DYRK1A and those without DYRK1A were affected to a similar extent (Figure 5D; Figure Supplementary 8A), suggesting that the effect of DYRK1A on RPG expression goes beyond its direct binding to promoters. The reduction in transcript levels of selected DYRK1A target and non-target RPGs was validated by RT-qPCR using two different shRNAs directed against DYRK1A (Figure Supplementary 7B and C). The analysis of the occupancy of Pol II at RPGs showed a general decrease of Pol II along RPGs gene bodies upon DYRK1A depletion, at both DYRK1A-positive and DYRK1A-negative RPGs (Figure 5E), pointing to a transcriptional effect as responsible for the reduction in RPG transcript steady-state levels.

All these results depict a complex scenario in which physiological levels of DYRK1A are important for maintaining RPG mRNA levels through at least two different, although not necessarily exclusive, mechanisms: on the one hand, through its recruitment to the proximal promoter regions of a subset of RPGs; and on the other hand, through an indirect mechanism that impacts on the levels of transcribing Pol II at the RPGs.

### DYRK1A depletion causes a reduction in ribosome content

We next wondered whether the downregulation of RPGs transcript levels induced by DYRK1A silencing would be reflected at the protein level. As such, we used mass spectrometry (MS) to quantify the ribosome composition in control cells and cells depleted in DYRK1A. A cytosolic fraction enriched in ribosomes was prepared from T98G cells by ultracentrifugation through a sucrose cushion and we analyzed two independent experiments with three biological replicates each, and considered for quantification only those proteins identified in at least three samples in any of the conditions. Our dataset had a strong overlap with those published in studies defining the human riboproteome (35, 36) (Figure Supplementary 9A). Indeed, not only were the biological functions related to protein synthesis (RPs and other well-known translation-associated proteins) among those enriched but also, those related to oxidative phosphorylation, RNA transport/processing or endocytosis (Figure Supplementary 9B; Supplementary Table S5), probably reflecting co-sedimenting complexes. An enrichment in nuclear proteins related with splicing has also been described (37). Nevertheless, core RPs represent more than 75% of the protein mass in the fraction analyzed (Figure 6A).

**Figure 6.**
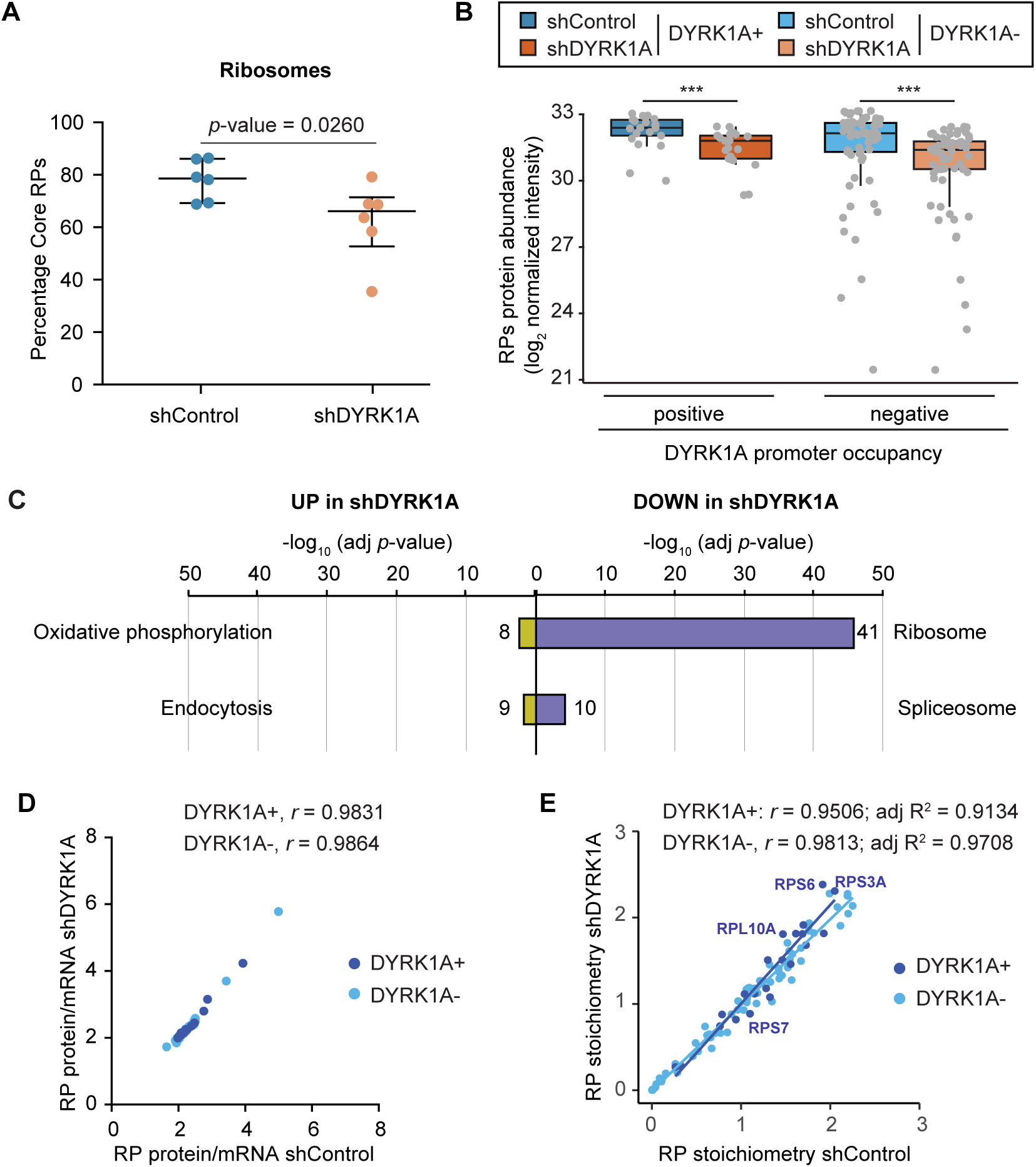
DYRK1A depletion causes a reduction in ribosome content. (**A**) The relative RP values (sum of all RP TOP3 values/sum all TOP3 values) for each sample of shControl (blue) and shDYRK1A (brown) T98G cells are represented, also showing the median and IQRs (n=6, two-tailed Mann-Whitney test). (**B**) Box plot indicating the levels of the RPs (log_2_ of normalized peptide intensities) separated into two clusters according to the presence of DYRK1A at the promoters of their corresponding genes, both in shDYRK1A and shControl T98G cells (Wilcoxon matched-pairs signed rank test: ***p<10^-4^). (**C**) Functional categorization of the proteins found more abundant in the shDYRK1A cells (UP) or in the shControl cells (DOWN) (proteins with p-value<0.05 in the comparisons or present in only one of the conditions were used). The number of proteins identified in each category is shown. (**D, E**) Correlation analysis of the ratio of RP protein and mRNA abundances (**D**) and RP stoichiometry (**E**) of shControl and shDYRK1A T98G cells. A color code was used to indicate the presence (+) / absence (-) of DYRK1A at the RPG promoter regions. The Spearman’s correlation coefficient is shown for each subset. In **E**, the value for the adjusted R^2^ for each set is also included.

The MS data allowed for the detection and quantification of all RPs except for the RPL10L and RPL39L paralogs, which showed low mRNA levels or not expressed at all in T98G cells, and for RPL41 that is usually not detected by MS approaches (35). In addition, this quantification also indicated a lower relative abundance of the RPS27L, RPL3L, RPL7L1, RPL22L1, RPL26L1 and RPL36AL paralogs than their corresponding pair (Figure Supplementary 10A and B; Supplementary Table 6), suggesting that they are under-represented in ribosomes from T98G cells. However, we cannot rule out that the variation in stoichiometry could be due to some RPs being loosely bound to the ribosome and thus, their presence may be affected by the method for cell extract preparation or because pools of RPs performing extra-ribosomal functions might not be present in the cell fraction analyzed. We observed variability in the protein/mRNA ratio in T98G cells, with extreme cases for most of the weakly expressed paralogs like RPL3L, RPL7L1 or RPL22L1 (Figure Supplementary 10A and B). Although we cannot rule out that some RPs are underrepresented in the fraction analyzed, the results are consistent with published data showing that the amounts of RPs correlate poorly with their corresponding mRNA levels in other cellular contexts (38). In agreement with the RNA data, our analysis did not reveal significant differences in the relative abundance of RPs encoded by DYRK1A-positive or DYRK1A-negative genes (Figure 6B; Figure Supplementary 10A and B).

We next focused on the alterations induced by silencing DYRK1A. The datasets revealed significant differences in the amount of protein associated with specific functional categories in the ribosomal-enriched fractions from control and DYRK1A depleted cells. Proteins in the oxidative phosphorylation category were enriched in the shDYRK1A cell fraction (Figure 6C), including components of the mitochondrial electron transport chain (COX6B1, COX7A2L, SDHB) or mitochondrial ATP synthases (Supplementary Table 6). Conversely, the ribosome category was enriched in the protein dataset reduced in the fraction from shDYRK1A cells (Figure 6C; Supplementary Table 6). Indeed, DYRK1A depletion significantly reduced the total ribosome mass when calculated relative to the total amount of protein in the fraction analyzed by MS (Figure 6A). In accordance with the general effect of DYRK1A on RPG transcript levels, less RP amounts were found for RPGs containing or lacking DYRK1A at promoters (Figure 6B; Figure Supplementary 10A and B). Notably, rRNA content tended to fall (Figure Supplementary 10D), supporting the decrease in ribosome amounts. The protein/mRNA ratios correlated strongly between control and DYRK1A silenced cells (Figure 6D), suggesting that changes in protein abundance were largely due to the changes in RNA abundance upon DYRK1A depletion. Finally, differences in RP stoichiometry were observed between shControl and shDYRK1A cells (Figure 6E), which might suggest differences in ribosome composition. All these results indicate that cells respond to DYRK1A depletion by reducing the steady state levels of RPs and that this effect occurs, at least in part, at the transcript level.

### DYRK1A plays a role in cell size control by regulating protein synthesis

Altered ribosome biogenesis can lead to major defects in translation and thus, we assessed the impact of DYRK1A on the translational status of cells. The functional status of ribosomes was first analyzed by polysome profiling on sucrose gradients upon DYRK1A silencing. Downregulation of DYRK1A diminished the polysome fractions, which corresponded to those ribosomes engaged in active translation, with a concomitant increase in the monosome peak (Figure 7A and B). These results indicate that DYRK1A depletion leads to polysome disorganization. Indeed, there was a significant reduction of translational rates in DYRK1A-depleted cells when measured through radiolabeled ^35^S-methionine incorporation (Figure 7C). This effect was specific, as it was clearly observed with two distinct shRNAs targeted to DYRK1A (Figure 7D).

**Figure 7.**
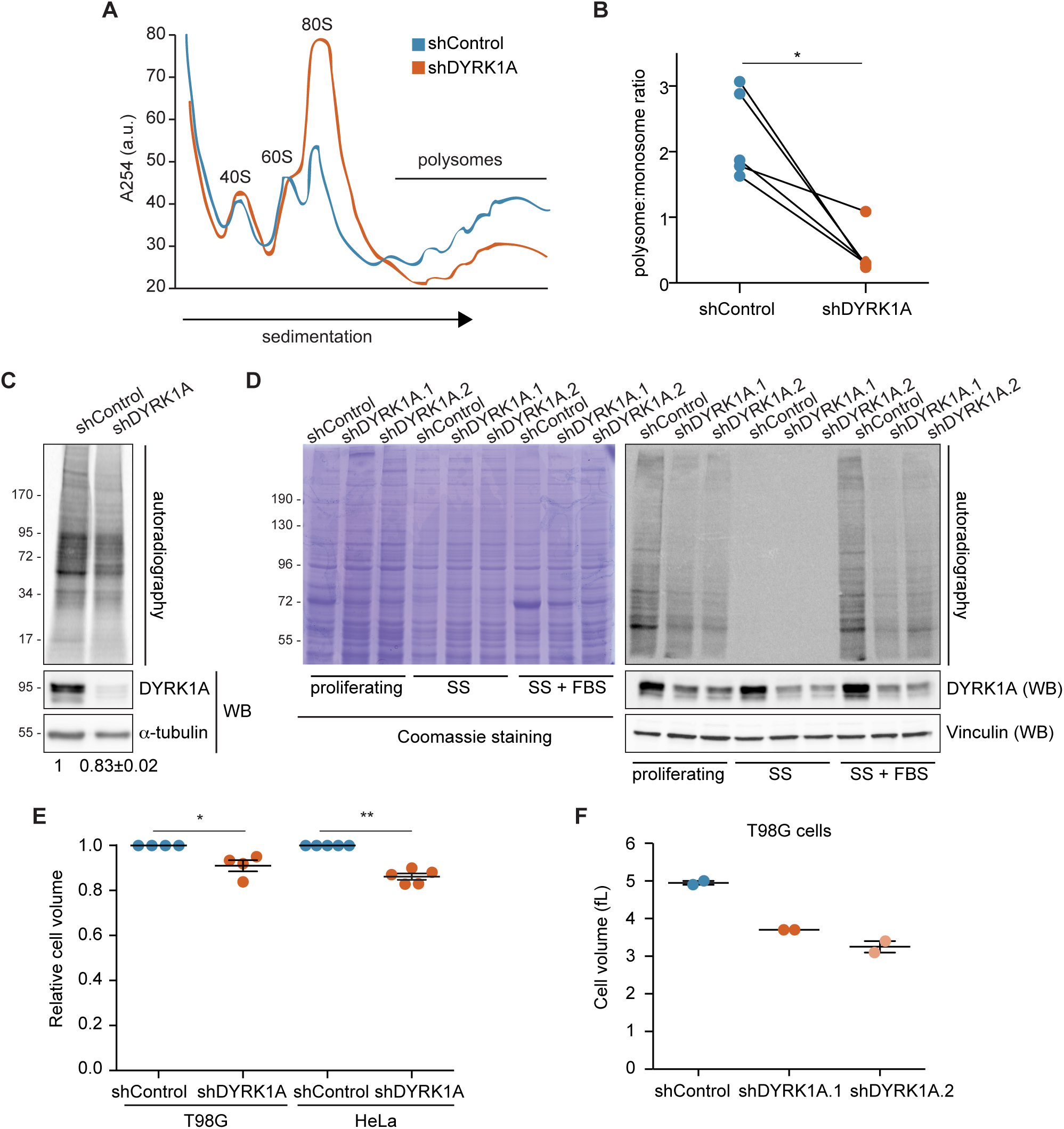
DYRK1A controls cell size by regulating protein synthesis. (**A**) Polysome profile obtained by sucrose gradient sedimentation of T98G cells in which DYRK1A is down-regulated via lentiviral delivery of shRNAs (shDYRK1A, orange line) and relative to the control cells (shControl, blue line). The position of the 40S, 60S, 80S and polysome peaks is indicated. The *y*-axis shows absorbance at 254 nm in arbitrary units and the *x*-axis corresponds to the different fractions, with the top of the gradient to the left. (**A**) The areas under the curve for polysomes and monosomes were measured from the polysome profiles of paired shControl and shDYRK1A experiments, and the polysome:monosome ratio was calculated (n=5, shDYRK1A.1 [3] and shDYRK1A.2 [2]; Wilcoxon matched-pairs signed ranks test: \**p*<0.05). (**C**) Global protein synthesis assays were performed by metabolic ^35^S-methionine labeling in shDYRK1A.1 or shControl T98G cells, visualizing the incorporation of the labeled amino acid by SDS-PAGE of the total cell extracts and autoradiography. The presence of DYRK1A was analyzed in WB using tubulin as a loading control. Quantification of the average radioactive intensity of independent experiments is shown at the bottom of the images, with the values of the shControl condition set as 1 (mean ± SEM, n=3, Wilcoxon matched-pairs signed ranks test, *p*<10^-3^). (**D**) Autoradiography of protein extracts prepared from proliferating, serum starved cells for 48 h (SS), or serum starved cells re-incubated with FBS for 30 min (SS+FBS) and pulse-labeled with ^35^S-Met for 20 min. Equal numbers of T98G cells infected with the indicated shRNA- lentivectors were used. The reduction in DYRK1A was assessed in WB using vinculin as a control and a Coomassie stained gel as a loading control is also shown. (**E**) Cell volume represented in arbitrary units with the control cells set as 1 (mean ± SEM; Mann-Whitney test, \**p*<0.05, \*\**p*<0.01). (**F**) Cell volume of T98G cells infected with lentivirus expressing two independent shRNAs against DYRK1A or a shRNA Control. The cell volume is represented in femtolitre (fL) and the average of two independent measurements is shown.

In eukaryotes, cell growth is coupled to cell cycle progression and therefore, global translation rates change during the cell cycle (39). Defects in the cell cycle have been associated with alterations in DYRK1A levels in different cellular environments (40). Indeed, we found that DYRK1A silencing alters the cell cycle balance in T98G cells, augmenting the population of cells in the G1 phase (Figure Supplementary 11A). Hence, we next checked whether the shift in cell cycle phases was associated with the reduced translation rates. As such, T98G cells were arrested in G1 by serum deprivation (Figure Supplementary 11B), which led to a strong reduction in the rate of translation (Figure 7D). Reversing serum depletion for 30 min induced protein synthesis (Figure 7D), although the cell cycle profiles were not changed (Figure Supplementary 11B). In these conditions, lower rates of translation persisted in cells with silenced DYRK1A relative to control cells (Figure 7D), suggesting that the reduction in protein synthesis upon DYRK1A silencing is independent of the alterations to the cell cycle.

As DYRK1A is a highly pleiotropic kinase, we wondered whether the reduction in translation could be due to an effect of DYRK1A on other signaling pathways that regulate protein synthesis. Therefore, we analyzed the effect of DYRK1A on one of the major signaling pathways that regulates protein translation, the mechanistic Target of Rapamycin (mTOR) pathway (41), assessing the Thr389 phosphorylation of the RPS6 kinase beta-1 (RPS6KB1 or p70S6K) that represents a late event in mTOR pathway activation. DYRK1A depletion did not alter Thr389-p70S6K phosphorylation (Figure Supplementary 12A and B) and likewise, the MS data showed no differences in the amount of RPS6 peptides phosphorylated at Ser235, Ser236 and Ser240 (Figure Supplementary 12C), targets of p70S6K downstream of mTOR (41). Accordingly, DYRK1A does not appear to affect the mTOR pathway under regular growth conditions. Other signaling pathways like cellular stress and unfolded protein response can inhibit protein synthesis through the Ser51 phosphorylation of the translation initiation factor eIF2α (42). However, the levels of Ser51-eIF2α phosphorylation remained unchanged in the absence of the DYRK1A (Figure Supplementary 13A and B), indicating that DYRK1A-dependent inhibition of translation is not mediated by eIF2α phosphorylation. Moreover, DYRK1A was not detected in the ribosome enriched fraction by MS or in immunoblots of polysome-associated fractions (Figure Supplementary 14), suggesting that it is not tightly bound to actively translating ribosomes and that it probably does not act directly on polysomes.

Finally, as reduced protein synthesis might affect cell mass, we checked whether cell size was affected by the loss of DYRK1A activity. Indeed, DYRK1A-silenced HeLa and T98G cells were both significantly smaller than their controls in terms of cell volume (Figure 7E and F). Hence, the fine-tuning of cellular DYRK1A levels is important to assure the proper size of human cells is maintained.

## DISCUSSION

In this study, we show that there is a subset of RPGs in mammals whose promoters are marked by the palindromic TCTCGCGAGA motif and the presence of the protein kinase DYRK1A as shown by chromatin recruitment analysis in different human and mouse cell lines. A motif enrichment analysis did not find the TCTCGCGAGA motif within RPG promoters in yeast, basal metazoa or plants (10), although a similar motif (CGCGGCGAGACC) was found within the proximal promoter regions of 28 RPGs in *Caenorhabditis elegans* (11). In *Drosophila*, no similar motifs were enriched in RPG promoters (9) but nevertheless, the DRE sequence has been proposed to mirror such a motif (33). By contrast, the TCTCGCGAGA motif is conserved in vertebrates and it is generally found in DNAse I accessible regions, leading to the proposal that it serves as a core promoter element in TATA-less promoters associated with CpG islands (31). These findings would suggest that the TCTCGCGAGA motif is a *cis*-regulatory element that arose later in evolution and that it might be linked to co-evolution with regulatory factors acting in *trans*. Besides DYRK1A, the transcriptional repressor ZBTB33 also uses the TCTCGCGAGA motif as a putative chromatin recruitment platform (32), and we confirm here its presence at RPG promoters that contain this motif. Whether DYRK1A and ZBTB33, or other proteins recruited to chromatin via the TCTCGCGAGA motif, compete or collaborate in the regulation of common target genes, including RPGs, is an issue that merits further exploration.

In *Drosophila*, RPG expression is regulated by a combination of two TFs, M1BP and DREF, which are associated with distinct subsets of RPGs through binding to their corresponding DNA sequence motifs. These two proteins are responsible for recruiting TRF2, which substitutes for TBP in the assembly of the PIC (12–14). Our analysis shows no evidence of such a regulatory network in humans. On the one hand, we observe TBP binding to almost all RPG promoters, regardless of whether or not they contain a TATA sequence, which indicates that TBP would be responsible for PIC assembly in human RPGs, as supported by the presence of the largest subunit of TFIID, TAF1. It is worth noting that the presence of DYRK1A at RPG promoters is correlated with more TBP binding, opening the question on the existence of a functional cross-talk between DYRK1A and TBP. TBPL1, a TRF2 homolog, also binds to almost all RPGs, consistent with the finding that TBPL1 is recruited to the PIC, not as a substitute for TBP but rather along with it in mouse testis (43). By contrast, we do not detect significant enrichment of the human M1BP and DREF homologs ZNF281 and ZBED1, at human RPG promoters. Therefore, a completely different set of regulators must exist in mammals to interpret, and to respond dynamically to growth and stress cues.

With the exception of the TCT motif, most of the sequences implicated in regulating the expression of RPGs in humans are only present in a subset of RPGs (12), including the binding motifs for the SP1, GABP, MYC and YY1 TFs. We have extended these results by analyzing the presence of these TFs at RPG promoter regions, revealing a more general distribution than that inferred through the presence of their binding sites. Therefore, no specific bias was found for the recruitment of TFs like SP1, YY1 or MYC to promoters containing the TCTCGCGAGA motif or positive for DYRK1A recruitment. By contrast, the RPGs that associate with DYRK1A are characterized by reduced GABP binding. The distinct distribution of DYRK1A and GABP could allow for differential regulation of subsets of RPGs in response to a variety of stimuli. Notably, both *DYRK1A* and *GABP* are located on human chromosome 21 and when in trisomy, their overexpression might contribute to the general increase in RPG mRNA transcripts in the brain of individuals with DS (Supplementary File 2).

Depletion of DYRK1A results in the downregulation of RPG transcripts, affecting both RPGs with DYRK1A bound at their promoters and RPGs devoid of DYRK1A. These results resemble those found with the manipulation of MYC levels resulting in changes in RPG transcripts not directly associated to the presence of MYC at the promoter regions of those genes (44, 45). In terms of RPGs that interact with DYRK1A, this reduction could be a direct effect of the loss of DYRK1A at their proximal promoters and the subsequent reduction of Pol II CTD phosphorylation, as shown for *RPS11* (19). However, additional mechanisms must exist to explain that the decrease in RPG mRNA levels occurs in a general manner. The reduction in Pol II occupancy at RPGs upon DYRK1A depletion would suggest that transcription might be a target. Even so, we cannot exclude that the reduction in DYRK1A levels induces post-transcriptional effects acting on RPG mRNAs steady-state levels, or alters the complex interactions between RPG mRNA synthesis and degradation and/or ribosome biogenesis that have been described in yeast (46, 47). Finally, the DYRK1A-dependent effect on increasing the population of cells in G1 could also be at play. Indeed, a reduction in RPG transcripts was detected in T98G cells when at G1 phase by serum starvation (Figure Supplementary 15). However, this reduction was increased in DYRK1A-silenced T98G cells (Figure Supplementary 15), suggesting that a DYRK1A-dependent mechanism is operating independently of the cell cycle.

Our results demonstrate that the production of RPs and ultimately, the number of ribosomes in proliferating cells, depends on the physiological levels of DYRK1A. This could be a direct consequence of the alterations in the RPG transcript levels since the protein/mRNA ratios are not significantly affected upon DYRK1A silencing. Additionally, yet not exclusive, compensating mechanisms at the level of ribosome biogenesis might also operate (1). In either case, the shortage of ribosomes upon a loss of DYRK1A provokes translational dysfunction, being cell size reduction one of the phenotypic outputs. The reduction in ribosomes could either globally affect translation or it may result in transcript-specific translational control. In this context, the pool of mRNAs associated with polysomes that respond differentially to DYRK1A downregulation still needs to be characterized. Together with the identification of *cis*-regulatory elements in these transcripts, this information will surely help to discriminate between the two possibilities and establish a mechanistic framework. Nonetheless, our findings do not rule out the existence of other DYRK1A-dependent effectors that contribute to altered ribosome biogenesis and/or translation, and further studies will help decipher how the effects of DYRK1A in the nucleus and cytoplasm converge to control cell growth. Noteworthy, such a multilayer regulatory effect is not uncommon in growth regulators. This is the case of the mTOR protein kinase that regulates ribosomal biogenesis by acting on the transcription of rRNA and ribosome protein components, directly on the basal machinery and through epigenetic regulation, and in addition, on translation at many different levels (41).

The physiological context for the activity of DYRK1A on translational control remains a matter of speculation at this stage. For instance, DYRK1A has been associated with the regulation of cell proliferation, and given that cell growth and proliferation are intimately linked (39, 48), this kinase could couple the cell cycle with protein synthesis by maintaining the amounts of RPs. In addition, DYRK1A plays essential roles in central nervous system development, not only influencing cell numbers but also their differentiation (20). Hence, the effect of DYRK1A on translation might contribute to its role as a regulator of neurite and axonal extension, events requiring post-mitotic cell growth. It is also possible that the effects of DYRK1A on cell mass may affect other tissues, particularly since heterozygous mouse models exhibit a global reduction in body size (49). In this regard, DYRK1A has been shown to phosphorylate Pol II at gene loci involved in myogenic differentiation (27), a process that requires increased protein synthetic rates (39). In a different context, defects in protein production are closely related to cancer since enhanced translation is required to boost cell proliferation (2, 3), and DYRK1A has both positive and negative effects on cell proliferation depending on the tumor context (21). Finally, mutations in specific RPGs produce very unique phenotypes (2), including craniofacial anomalies and urogenital malformations (50). In addition, *RPL10* mutations have been linked to neurodevelopmental conditions including autism spectrum disorders and microcephaly (51), and indeed, translation is a process targeted in autism-associated disorders (52). All these features are hallmarks of *DYRK1A* haploinsufficiency syndrome in humans or of animal models with *Dyrk1a* dysregulation (24).

In summary, our findings have uncovered DYRK1A as a novel player in the regulation of translation and cell growth, further expanding the complex biology of this protein kinase and opening novel avenues for future work.

## MATERIAL AND METHODS

### Cell culture and lentivirus-mediated transduction

The HEK-293T, HeLa, U2OS and T98G cell lines were obtained from the American Type Culture Collection (www.atcc.org). E14 (129/Ola) mESCs were kindly provided by Maria Pia Cosma’s laboratory. Human cells were cultured in Dulbecco’s Modified Eagle’s Medium (DMEM) supplemented with 10% (v/v) fetal bovine serum (FBS) and antibiotics (100 U/ml penicillin and 100 μg/ml streptomycin). mESCs were cultured in 0.1% gelatin coated plates, with DMEM supplemented with 15% FBS, 2 mM L–glutamine, 1 mM sodium pyruvate, 0.1 mM non-essential amino acids, 0.5 mM 2-mercaptoethanol, 1000 U/ml ESGRO recombinant mouse Leukemia Inhibitory Factor and antibiotics. For serum starvation, T98G cells were grown in DMEM without FBS for 16-48 h. All cell lines were grown at 37 °C and in a 5% CO_2_ atmosphere.

Lentiviral stocks were generated by transfecting HEK-293T cells by the calcium phosphate precipitation method with pCMV-VSV-G envelope plasmid, pCMV-dR8.91 packaging construct, and pLKO-based Sigma Mission plasmids expressing shRNAs, either a non-targeting vector (shControl) or two shRNAs directed against DYRKA: DYRK1A.1 and DYRK1A.2. Lentivirus-containing supernatants were harvested 48 h and 72 h after transfection, and concentrated by ultracentrifugation (87,500 *xg*, 2 h, 4 °C). For cell transduction, the virus was added to the medium with the cells in suspension, and replaced with fresh media 24 h after infection and seeding. Infected cells were selected by adding 1.25 μg/ml of puromycin for 48-72 h, and the cells were allowed to recover in the absence of puromycin for 24 h.

### Cell cycle profile and volume determination

T98G cell pellets were fixed by dropwise addition of cold 70% ethanol and their DNA was stained with 4’,6-diamino-2-phenylindole (1 μg/ml, Roche), 0.1% Triton X-100 in phosphate-buffered saline (PBS). Cells were analyzed with a LSR II flow cytometer (Becton Dickinson, Franklin Lakes, New Jersey) with FACSDiva^TM^ software v6.1.2. Cell volume was determined with the Beckman Coulter Z2 Cell and Particle Counter. Cells were harvested by trypsinization, washed in cold PBS and measured in triplicate. Cell-type dependent cursor settings were used to discriminate between vital and dead cells/cell debris.

### Protein synthesis determination

To analyze global protein synthesis, T98G cells were incubated for 90 min in Methionine-free DMEM with 10% dialyzed FBS, metabolically labeled for 20 min with ^35^S-Met (50 μCi ^35^S-Met, 1175 Ci/mmol, Perkin Elmer) and then lysed in SDS-lysis buffer (25 mM Tris-HCl pH 7.5, 1% sodium dodecyl sulfate [SDS], 1 mM ethylenediaminetetraacetic acid [EDTA], 10 mM Na_4_P_2_O_7_, 20 mM β-glycerol phosphate). The protein extracts were resolved by SDS-PAGE and the incorporation of ^35^S-Methionine was detected by autoradiography of the dried gel using film or a Phosphoimager (Typhoon Trio, GE Healthcare, Freiburg, Germany).

### Preparation of polysome and ribosome enriched fractions

Polysome profiles were obtained from approximately 1×10^7^ T98G cells. Protein synthesis was arrested by incubation with cycloheximide (CHX, 100 μg/ml). The cells were washed in PBS containing CHX (100 μg/ml), collected in 1 ml of polysome lysis buffer (10 mM Tris-HCl pH 7.4, 100 mM NaCl, 10 mM MgCl_2_, 1% Triton X-100, 20 mM dithiothreitol [DTT], 100 μg/ml CHX, 0.25% sodium deoxycholate) and frozen rapidly in liquid nitrogen. Cell debris and nuclei were eliminated by centrifugation (12,000 *xg*, 5 min, 4 °C) and the nucleic acid concentration in the supernatants was assessed by measuring the A_260_ in a NanoDrop™ (Thermo Fisher Inc.). Samples with A_260_ ≈10 were loaded onto a 10-50% linear sucrose gradient prepared in polysome gradient buffer (20 mM Tris-HCl pH 7.4, 100 mM NH_4_Cl, 10 mM MgCl_2_, 0.5 mM DTT, 100 μg/ml CHX), and centrifuged in a Beckman SW41Ti rotor (35,000 rpm, 3 h, 4 °C). Profiles were obtained by continuous monitoring of the A_254_ (Econo-UV Monitor and Econo-Recorder model 1327; Bio-Rad Laboratories Inc., Barcelona, Spain). To calculate the polysome:monosome ratio, the polysome and monosome area under the curve was measured with ImageJ (1.50i) (53).

To isolate the total ribosome fraction, cells were collected in sucrose buffer (250 mM sucrose, 250 mM KCl, 5 mM MgCl_2_, 50 mM Tris-HCl pH 7.4, 0.7% Nonidet P-40), the cytosol was isolated by centrifugation (750 x*g*, 10 min, 4 °C), and then centrifuged again to obtain a post-mitochondrial supernatant (12,500 x*g*, 10 min, 4 °C). The supernatant was adjusted to 0.5 M KCl, and the volume equivalent to OD_260_=5 was loaded onto a sucrose cushion (1 M sucrose, 0.5 M KCl, 5 mM MgCl_2_, 50 mM Tris-HCl pH 7.4) and centrifuged in a Beckman TLA 100.3 rotor (250,000 x*g*, 2 h, 4 °C).

### Western blotting

To prepare the total cell lysates, cells were resuspended in SDS-lysis buffer and heated for 10 min at 98 °C. The protein extracted was quantified with the BCA Protein Assay Kit. For Western blotting, cell lysates were resolved on SDS-polyacrylamide gels (SDS-PAGE) and transferred onto Hybond-ECL nitrocellulose membranes. The membranes were blocked for 1 h at room temperature with 10% non-fat milk diluted in TBS-T (10 mM Tris-HCl pH 7.5, 100 mM NaCl, 0.1% Tween-20) and then incubated for 16 h at 4 °C with the primary antibodies diluted in TBS-T containing 5% non-fat milk (or bovine serum albumin [BSA] for the phosphospecific antibodies). After several washes in TBS-T, the membranes were incubated with horseradish peroxidase-conjugated secondary antibodies (Dako, Agilent Technologies, Santa Clara, California) for 1 h at room temperature diluted in TBS-T containing 5% non-fat milk. After washing in TBS-T, the chemiluminiscence signal was revealed with Western Lightning® Plus ECL and measured in a LAS-3000 image analyzer (Fuji PhotoFilm, Tokyo, Japan) with the LAS3000-Pro software. Band intensities were quantified with the ImageQuant^TM^ TL software or ImageStudio^TM^ Lite, and the relative protein levels were calculated using α-tubulin or vinculin as loading controls. Primary antibodies used are listed in Key Resources table.

### Mass spectrometry (MS) analysis

#### Sample preparation

Ribosome-enriched pellets were washed three times with 100 mM ammonium bicarbonate (ABC) and resuspended in 6 M Urea-100 mM ABC. Extracts (10□g) were incubated for 1 h at 37 °C in the presence of 0.3 mM DTT, followed by incubation with 0.6 mM iodoacetamide (Sigma) for 30 min at room temperature in the dark. Proteins were digested for 16 h at 37 °C with Lys-C endoprotease (1□g; Wako, Fujifilm) and then incubated for 8 h at 37 °C with sequence-grade trypsin (1□g; Promega, Madison, Wisconsin). Peptides were desalted on Ultra Micro Spin Columns C18 (The Nest Group Inc., Ipswich, Massachusetts) (54) prior to liquid chromatography (LC)-MS/MS.

#### Chromatographic and MS analysis

Samples were analyzed on a LTQ-Orbitrap Fusion Lumos mass spectrometer (Thermo Fisher Scientific) coupled to an Easy-nLC 1000 (Thermo Scientific, Proxeon) at the CRG/UPF Proteomics Unit. Peptides were loaded onto the analytical column and separated by reversed-phase chromatography on a 50 cm column with an inner diameter of 75□ m, packed with 2□ m C18 particles (Thermo Scientific). Chromatographic gradients started at 95% buffer A (0.1% formic acid in water) and 5% buffer B (0.1% formic acid in acetonitrile) with a flow rate of 300 nl/min for 5 min, and they gradually increased to 78% buffer A - 22% buffer B in 79 min, and then to 65% buffer A - 35% buffer B in 11 min. After each analysis, the column was washed with 10% buffer A - 90% buffer B for 10 min. The mass spectrometer was operated in positive ionization mode with nanospray voltage set at 2.4 kV and the source temperature at 275 °C. Ultramark 1621 was used for external calibration of the FT mass analyzer and an internal calibration was performed using the background polysiloxane ion signal at m/z 445.1200. Acquisition was performed in data dependent acquisition (DDA) mode and full MS scans with 1 micro scans were used at a resolution of 120,000 over a mass range of m/z 350-1500. Auto gain control (AGC) was set to 1E5 and charge state filtering that disqualifies single charged peptides was activated. In each cycle of DDA analysis, the most intense ions above a threshold ion count of 10,000 were selected for fragmentation following each survey scan. The number of selected precursor ions for fragmentation was determined by the “Top Speed” acquisition algorithm with a dynamic exclusion of 60 s. Fragment ion spectra were produced via high-energy collision dissociation at a normalized collision energy of 28% and they were acquired in the ion trap mass analyzer. AGC was set to 1E4, using an isolation window of 1.6 m/z and a maximum injection time of 200 ms. All data were acquired with Xcalibur software (v4.1.31.9, Thermo Fisher Scientific). Digested BSA was analyzed between each sample to avoid sample carry over and to assure the stability of the instrument. QCloud (55) was used to control the instrument’s longitudinal performance. The spectra acquired were analyzed with the Proteome Discoverer software suite (v2.3, Thermo Fisher Scientific) and the Mascot search engine (v2.6, Matrix Science Ltd., London, UK) (56). The data was used to search the SwissProt human database (February 2020), including a list of common contaminants and the corresponding decoy entries (57).

For peptide identification, a precursor ion mass tolerance of 7 ppm was used for MS1, with trypsin as the chosen enzyme and up to three miscleavages allowed. The fragment ion mass tolerance was set to 0.5 Da for MS2. Oxidation of methionine and N-terminal protein acetylation were used as variable modifications, whereas carbamidomethylation on cysteine was set as a fixed modification. In the analysis of phosphorylated peptides, phosphorylation of serine, threonine and tyrosine were also set as variable modifications. False discovery rate (FDR) was set to a maximum of 5% in peptide identification. Protein quantification was retrieved from the protein TOP3 Area node from Proteome Discoverer (v2.3). For normalization, a correction factor was applied: sum TOP3 replicate “n”/average sum TOP3 all replicates. Based on the estimated normalized abundance, values were log_2_ transformed, and the fold change (FC) and *p*-values were calculated. Two independent experiments, each with three biological replicates, were performed on T98G cells transduced with shControl or shDYRK1A lentiviruses, and only those proteins detected in at least three replicates of any condition were quantified. To study RP stoichiometry, the intensity of each RP was defined relative to the intensities for all RPs. The RP protein/mRNA ratios were obtained using the log_2_ transformed normalized protein abundance and the log_2_ transformed normalized RNA counts from the RNA-Seq experiments.

### Chromatin immunoprecipitation (ChIP)

#### Sample preparation

Formaldehyde-crosslinked cells were washed twice with cold PBS, resuspended in 1 ml of Lysis buffer I (5 mM PIPES pH 8.0, 85 mM KCl, 0.5% NP-40, protease inhibitor cocktail - cOmplete Mini) and incubated on ice for 10 min. After centrifugation (800 *xg*, 5 min, 4 °C), the cell pellets were resuspended in 0.6 ml of Lysis buffer II (1% SDS, 10 mM EDTA, 50 mM Tris-HCl pH 8.0, protease inhibitor cocktail) and incubated for additional 10 min on ice. Chromatin was sonicated to an average size of 0.2-0.5 kb with a Bioruptor (Diagenode SA, Seraing, Belgium) and the chromatin corresponding to 150 μg DNA was diluted in 1 ml of Buffer III (300 mM NaCl, 200 mM Tris-HCl [pH 8], 10 mM EDTA, 1% SDS, 5% Triton X-100) and the samples were incubated with specific antibodies or control rabbit IgGs (listed in Key Resources Table) for 16 h at 4 °C with rotation. The immunocomplexes were recovered by incubating with 30 μl of Protein A-Sepharose beads for 3 h at 4 °C with rotation, and the beads were then washed three times with low salt buffer (50 mM HEPES pH 7.4, 140 mM NaCl, 1% Triton X-100), once with high salt buffer (50 mM HEPES pH 7.4, 500 mM NaCl, 1% Triton X-100), once with LiCl buffer (10 mM Tris-HCl pH 8, 250 mM LiCl, 1% NP-40, 1% deoxycholic acid, 1 mM EDTA), and once with TE buffer (100 mM Tris-HCl pH 7.5, 50 mM EDTA). All wash buffers were supplemented with the protease inhibitor cocktail. DNA was eluted by incubation in elution buffer (1% SDS, 100 mM NaHCO_3_) in two steps of 30 min, each at 65 °C, and crosslinking was reverted by an additional incubation in 200 mM NaCl for 16 h at 65 °C. Finally, chromatin-associated proteins were degraded by adding Proteinase K (1.6 U) for 2 h at 45 °C, and the DNA was purified by phenol/chloroform extraction and ethanol-precipitation. The DNA was quantified with the Qubit→ dsDNA HS Assay Kit.

#### Library generation and sequencing

DNA libraries were generated with the Ovation→ Ultralow Library System V2. Libraries were sequenced with 50 bp single end reads on an Illumina Hiseq-2500 sequencer at the CRG Genomics Unit. The quality of the sequenced reads was controlled using with FastQC (58). Reads were mapped to the human or mouse genome assemblies (hg38 and mm9, respectively) using Bowtie (59), and *p*-values for the significance of ChIP-Seq counts were calculated relative to the input DNA (60), using a threshold of 10^-8^ and FDR <1%. To analyze RPG promoter occupancy, datasets from The Encyclopedia of DNA Elements (ENCODE) Consortium were used (61, 62) (https://www.encodeproject.org/): GABPA: dataset GSE32465 (sample GSM1010739); MYC: dataset GSE33213 (sample GSM822290); SP1: dataset GSE32465 (sample GSM803363); TAF1: dataset ENCSR000BKS (sample ENCFF101GBL); TBP: dataset GSE31477 (samples GSM935606, GSM935277, GSM935495); TBPL1: dataset ENCSR783EPA (sample ENCFF167PVP); YY1: dataset GSE32465 (samples GSM803406, GSM1010897); ZBED1: dataset ENCSR286PCG (sample ENCFF465PNK); ZNF281: dataset ENCSR403MJY (sample ENCFF948PYK). Read numbers were normalized to a 1x reads per million (RPM) sequencing depth in both this work and the ENCODE datasets.

### RNA-Seq

#### Sample preparation

RNA was isolated with the RNeasy extraction kit and the samples were treated with DNase I (2 U/μl:) for 30 min at 37 °C. RNA was quantified with NanoDrop and quality-controlled on a Bioanalyzer 2100 (Agilent). For T98G cell spike-in normalization, equal numbers of T98G cells for each condition were mixed with a fixed number of *D. melanogaster* Kc167 cells (1:4 ratio). For reverse transcription (RT), 0.5-1 μg RNA was subjected to cDNA synthesis with Superscript II Reverse Transcriptase and random primers.

#### Library generation and sequencing

Libraries were prepared with the TruSeq Stranded mRNA Sample Prep Kit v2 and sequenced with Illumina Hiseq-2500 to obtain 125 bp pair-ended reads.

#### Data analysis

Genomic reads were mapped and counted using STAR (63), plotting T98G cells against the human and fly genomes (hg38 and dm3, respectively) or HeLa cells against the human genome alone. Those reads that could not be uniquely mapped to just one region were discarded. Differential gene expression was assessed with the DESeq2 package in R, filtering genes that had >10 average normalized counts per million (64). For the spike-in libraries, the size factor of each replicate was calculated according to exogenous *Drosophila* spike-in reads. Expression was considered to be altered when *p*-value≤0.05, and with a log_2_(FC) above 0.7 and below -0.7 for up- and down-regulated genes, respectively. To measure RPG expression level in U2OS cells, we analyzed the GSE117212 dataset in the same manner (GSM3274834, GSM3274835, GSM3274837) (65), obtained from the NCBI’s Gene Expression Omnibus (GEO) repository (https://www.ncbi.nlm.nih.gov/geo/) (66). To analyze the expression of RPGs in Down syndrome individuals, the GEO dataset GSE5390 (67) was analyzed with GEO2R using the default parameters.

### Quantitative PCR (qPCR)

PCR reactions were performed in triplicate in 384 well plates with SYBR Green (Roche) and specific primers, using a Roche LC-480 machine. The crossing point was calculated for each sample with the Lightcycler 480 1.2 software. No PCR products were observed in the absence of template and all primer sets gave narrow single melting point curves. For ChIP-qPCR, a 1/10 dilution of ChIP DNA was used as the template for the PCR reaction, and a 1/1000 dilution of input DNA was used as standard for normalization. For RT-qPCR, 1/10 dilution of the cDNAs was used and expression of the *D. melanogaster* gene *Act42A* was used for normalization.

### Computational tools

The Integrated Genomics Browser v9.0.0 (68) was used to visualize the ChIP-Seq data sets and an in-house gene and promoter annotation pipeline was used for peak annotation. The script uses data from a consensus position weight matrix (PWM) for the presence of the DYRK1A motif (19), and the University of California Santa Cruz (UCSC) genome browser software (http://genome.ucsc.edu/) (69) via the UCSC Table Browser (70) for Ensembl/NCBI gene and transcript annotation. Extra annotations were provided using Biomart (71). BEDTools (72) was used to create overlaps of significant peaks with genomic annotated regions with the “BEDtools intersect” command. Density plots and heatmaps to profile the average binding of the factors analyzed on defined genomic intervals were generated with computeMatrix and plotHeatmap tools from deepTools v3.0.0 (73). Four genes (*RPL10L*, *RPL3L*, *RPS18* and *RPS4Y2*) were excluded from the analysis because they were not bound by any of the factors. The DYRK1A-PWM (19) was used to define RPG promoters containing the DYRK1A consensus motif within a -1000 bp to +100 bp region from TSS using the FIMO package (74) with *p*-values≤ 3×10^-4^. The *p*-value was converted to a *q*-value following the method of Benjamini and Hochberg (75). *Bona-fide* motifs were considered when *p*-value < 10^-4^ and degenerated motifs were those with 10^-4^ < *p*-value < 3×10^-4^. CentriMo (30) was used to identify the significant preference of the DYRK1A motif for particular locations within DYRK1A binding RPGs over a ±250 bp region of ChIP-Seq peaks summit. To identify RPG promoters harboring the TCT-motif (12), sequences from -50 to +50 (relative to the C+1 transcription start site) were retrieved from the Eukaryotic Promoter Database EPD (https://epd.epfl.ch//index.php; (76) and analyzed using the MEME tool in the MEME Suite (http://meme-suite.org) (77). The Jaspar database was used to retrieve known transcription factor binding profiles (http://jaspar.genereg.net/) (78)). The Jaspar-associated Myc-PWM MA0147.3 was used to define RPG promoters containing the Myc motif within a - 1000 bp to +100 bp region from TSS using the FIMO package (74) with *p*-values≤1×10^-4^. The Enrichr webtool (79) was used to identify pathways enriched in the proteomics datasets (KEGG 2019 Human). The BioVenn free software (80) was used to overlap gene/protein targets from different datasets. Human orthologs of the *Drosophila* proteins were found in FlyBase (release FB2018_06; https://flybase.org/) (81).

Scatter and box plots were generated with the R package ggplot2 (82). For the box plots, the bottom and top of the box are the first and third quartiles, and the band inside the box is the median. The ends of the whiskers represent the lowest data point still within a 1.5 interquartile range (IQR) of the lower quartile, and the highest data point still within 1.5 IQR of the upper quartile. Any data not included between the whiskers is plotted as an outlier with a dot. Bar graphs were generated with Microsoft Excel v15.33.

### Statistical analysis

To calculate statistical significance, the normality of the samples was evaluated with the Shapiro-Wilk normality test (Prism 5, v5.0d), and parametric or non-parametric tests were used accordingly. Statistical significance was calculated with two-tailed Mann-Whitney or Student’s tests for unpaired samples or with a Wilcoxon matched pairs signed ranks test (Prism 5, v5.0d), and *p*-value≤0.05 considered as significant. All experiments were performed independently at least three times.

## ACKNOWLEDGEMENTS

We thank all the members of Susana de la Luna’s laboratory for their helpful discussions, Alicia Raya for technical assistance, the members of Juana Díez’s laboratory (Molecular Virology lab, UPF) for assistance with the polysome profiling, Enrique Blanco (Epigenetics Events in Cancer Lab, CRG) for helpful discussions regarding the RNA-Seq analysis, Sarah Bonnin (CRG Bioinformatics Unit) for advice on the bioinformatics analysis, Aránzazu Rosado (Genome architecture Lab, CRG) for kindly providing the Kc167 *Drosophila* cells, Giorgio Dieci (Dept. of Chemistry, Life Sciences and Environmental Sustainability, University of Parma) for critical reading of the manuscript, and Mark Sefton for English language editing. We acknowledge the assistance of Eduard Sabidó and Eva Borras from the CRG/UPF Proteomics Unit and support from the CRG Genomics Facility.

## DATA AVAILABILITY

All the raw and processed sequencing data generated in this study have been submitted to the NCBI GEO repository under accession number GSE155809. The raw proteomics data have been submitted to the Proteomics Identifications Database (PRIDE) under accession number PXD022966.

## AUTHOR CONTRIBUTIONS

S.d.l.L. supervised the project and acquired funding. C.d.V., L.B. and S.d.l.L. designed the experiments. C.d.V. and L.B. acquired the data. C.d.V., L.B., R.F. and S.d.l.L. analyzed the data. C.d.V. performed high throughput sequencing data analysis. S.d.l.L. wrote the manuscript together with C.d.V. All authors contributed critically and approved the final manuscript.

