## Supplementary Figures for "Loss of the DYRK1A protein kinase results in reduction of ribosomal protein genes expression, ribosome mass and translation defects"

#### SUPPLEMENTARY FIGURE LEGENDS

**Figure S1. Supporting data for the specificity of the DYRK1A antibody in the ChIP-seq experiments shown in Figure 1.**

(**A**) Metagene plot showing DYRK1A occupancy of human RPGs in shControl (blue) or shDYRK1A (orange) infected T98G cells. The *y*-axis represents the relative protein recruitment, quantified as significant (sig) ChIP-Seq tags. (**B**) Representative example of DYRK1A occupancy at a DYRK1A binding (*RPL27A*) or a DYRK1A-negative (*RPS5*) RPG in T98G cells infected with the shControl or shDYRK1A lentivirus. The arrow indicates the direction of transcription.

**Figure S2. The transcription factor ZBTB33 is recruited to RPG promoters positive for DYRK1A and the TCTCGCGAGA DNA-motif.**

(**A**) Density plots showing DYRK1A and ZBTB33 ChIP binding relative to the TSS of all human RPGs. The heatmaps show DYRK1A and ZBTB33 occupancy across all human RPGs relative to the TSS in T98G cells. The binding score (-$\log_{10}$ Poisson *p*-value) is indicated by the color scale bar and the offset was ± 3 kb from the TSS. (**B**) Representative examples of RPGs occupied by DYRK1A and ZBTB33. The arrows indicate the direction of transcription. (**C**) CentriMo plot showing the distribution of the DYRK1A motif in the RPG promoters that bind ZBTB33. The solid curve shows the positional distribution of the best strong DYRK1A motif site at each position in the ZBTB33 RPG-ChIP-Seq peaks (500 bp). The *p*-value for the central motif enrichment is indicated.

**Figure S3. Supporting data for the recruitment of ZBED1 to human RPG promoters positive for DYRK1A shown in Figure 3.**

(**A**) CentriMo plot showing the distribution of the DYRK1A motif for the RPG promoters that bind ZBED1. The solid curve shows the positional distribution (averaged over bins 10 bp in width) of the best strong site of the DYRK1A motif at each position in the RPG-ChIP-Seq peaks (500 bp). The *p*-value for the central motif enrichment is indicated. (**B**) Density plot of the ZBED1 occupancy at bound RPGs (42 out of 90 RPGs) relative to center of the ZBED1 peaks. ZBED1-bound RPGs were classified as either DYRK1A positive or negative RPGs relative to center of the ZBED1 peak.

**Figure S4. Recruitment of MYC, SP1 and YY1 to RPG promoters in comparison with that of DYRK1A.**

(**A**) The heatmaps show MYC, SP1 and YY1 (GM12868 cells, ENCODE datasets GSE33213 and GSE32465) occupancy on human RPGs. The genomic region considered is shown on the *x*-axis and the color scale bar indicates the binding score. (**B**) Unbiased *k*-mean clustering of the average binding of DYRK1A (T98G cells, this work), MYC, SP1 and YY1 (GM12868 cells, ENCODE datasets GSE33213 and GSE32465) to all RPGs. The color scale bar indicates binding score and the genomic region considered is shown on the *x*-axis.

**Figure S5. RPG mRNA expression in different human cell lines.**

(**A-C**) The heatmaps correspond to the top 5% more expressed genes in T98G (**A**), HeLa (**B**) and U2OS (**C**) cells. RPGs are highlighted on the right-hand side in each panel. The color bar in panel **A** represents normalized read counts from RNA-Seq experiments (T98G and HeLa, this work; U2OS, ENCODE dataset GSE117212). (**D-F**) Box plots for the expression of genes corresponding to the 5% of the most strongly expressed genes, with RPGs removed (no-RPGs) or only considering RPGs (RPGs) in T98G (**D**), HeLa (**E**) and U2OS (**F**) cells. The number of genes analyzed is indicated in parenthesis above each graph.

**Figure S6. Expression analysis of RPGs in different cell lines according to the presence of DYRK1A at their promoter regions.**

Box plot indicating RPG transcript levels (normalized counts) separated into two clusters depending on the presence of DYRK1A at their promoters based on the RNA-Seq data from

HeLa cells (**A**, this work) or U2OS cells (**B**, GEO dataset GSE117212) (unpaired Mann-Whitney test, ns=not significant).

**Figure S7. Global alterations in gene expression upon DYRK1A depletion.**

(**A**) Differentially expressed genes in T98G cells upon DYRK1A silencing as analyzed by RNA-Seq after spike-in normalization (adj *p*-value < 0.05; DOWN, $\log_2$(FC) < -0.7; UP, $\log_2$(FC) > 0.7; ns, not significant). Dotted lines indicate the thresholds of statistical significance. (**B**) Overlap between differentially expressed genes (DEG) as determined by RNA-Seq and genes with DYRK1A at their promoters witnessed by ChIP-Seq in T98G cells.

**Figure S8. Supporting data for the alterations in RPG transcript levels in DYRK1A depleted cells shown in Figure 5.**

(**A**) The graph presents the $\log_2$(FC) values for RPG mRNA transcripts with an adj *p*-value < 0.05 clusterized according to DYRK1A presence at their promoters in shDYRK1A versus shControl comparisons (two-tailed Mann Whitney test, ns=not significant). (**B**) RT-qPCR validation of differentially expressed RPGs in T98G cells transduced with shControl or shDYRK1A lentiviruses. DYRK1A and one translation-related non-RPG factor (*EIF4E2*) RNA levels are also shown. The data was corrected by *Drosophila* spike in values and represented as the RNA levels relative to the shControl arbitrarily set as 1 (mean ± SD of independent experiments: Mann-Whitney test, *$p\leq0.05$, **$p\leq0.01$, ***$p\leq0.001$, ns=not significant). (**C**) RT-qPCR validation of the expression of selected RPGs in T98G cells upon DYRK1A downregulation following lentiviral transduction of two distinct shRNAs (shDYRK1A.1 and shDYRK1A.2). The data are represented as the RNA levels relative to the shControl condition set arbitrarily as 1 (mean ± SD of 3 technical replicates) and the reduction in DYRK1A levels is shown in WBs. (**D**) The levels of DYRK1A were assessed in WBs of samples corresponding to the experiment shown in Figure 5D.

**Figure S9. Supporting data for the riboproteome analysis shown in Figure 6.**

(**A**) Venn diagram indicating the overlap between the riboproteomes identified in Reschke et al (2013) and Imami et al (2018), and the proteins identified by MS in this work. (**B**) Functional categorization of the proteins identified by MS, showing the number of proteins identified in each category.

**Figure S10. RP expression analysis based on MS quantitative data in T98G cells comparing shControl and shDYRK1A.**

(**A, B**) Relative abundance of RPs from the large (**A**) and small (**B**) ribosomal subunits in shControl and shDYRK1A T98G cells. The normalized level of each RP is shown relative to the mean of all RP values (n=6, for each condition; unpaired, two-tailed, Mann-Whitney test: *$p$<0.05; **$p$<0.001). RPs whose encoding genes present DYRK1A at the promoter level are depicted in bold. (**C**) RPG mRNA levels determined by RNA-Seq are plotted relative to the normalized protein levels obtained by MS quantification. Both values are represented as $\log_2$ values, and the blue and light blue dots correspond to RPGs bound by DYRK1A or not, respectively. (**D**) Quantification of 18S and 28S rRNAs was obtained by Bioanalyzer calculation of the areas of each of the peaks. The graph represents the mean ±SEM of 3 independent RNA samples for each condition (ns, not significant with and unpaired, two-tailed, Mann-Whitney test, yet differences for the 18S rRNA were significant when using the Student's test).

**Figure S11. Cell cycle profile analysis of T98G cells depleted of DYRK1A or under serum starvation.**

(**A**) The cell cycle profiles of T98G shControl or shDYRK1A infected cells were determined by FACs analysis, and the distribution of the G1, S and G2/M subpopulations is shown (mean ± SD, n=3 biological replicates, two-way Anova for multiple comparisons: ***$p$≤0.001, ns=not significant). Note that depletion of DYRK1A leads to an increase in the cells in G1 phase, concomitant with a reduction of cells in S phase. (**B**) Cell cycle profile of T98G cells grown under normal growth conditions (proliferating) and after serum starvation (SS) to show the increase in the G1 population. Re-addition of FBS (SS +FBS) did not induce changes in the cell cycle profile at 2 h after serum addition.

**Figure S12. Analysis of the alterations in p70S6K phosphorylation levels in response to DYRK1A silencing in T98G.**

Analysis of the Thr389 phosphorylation of p70S6K (p-p70S6K) assessed in WB of shControl or shDYRK1A T98G cells. A representative experiment (**A**) and the quantification of the p-p70S6K signal relative to the total p70S6K (**B**) are shown (mean ± SD, n = 4, Wilcoxon matched-pairs signed ranks test, ns=not significant). The dots and lines show independent experiments. (**C**) TOP3 quantification of the RPS6 peptide containing phosphorylated Ser236 and Ser240 from shControl and shDYRK1A samples in MS experiments. RPS6 quantification in each sample is also shown, as well as the ratios between the two values.

**Figure S13. Analysis of the alterations in eIF2α phosphorylation levels in response to DYRK1A silencing in T98G.**

Analysis of eIF2α Ser51 phosphorylation (p-eIF2α) in extracts from shControl or shDYRK1A T98G cells. A representative WB (**A**, *non-specific band) and the quantification of the phospho-eIF2α to total eIF2α ratio (**B**) is shown (mean ± SD, n =3: Wilcoxon test, ns = not significant). The dots and lines show independent experiments.

**Figure S14. DYRK1A does not associate with a polysome fraction.**

WB analysis of fractions from a polysome gradient to detect the presence of DYRK1A. Vertical lines indicate that the panel is not continuous. In the graph, the *y*-axis shows absorbance at 254 nm in arbitrary units and the *x*-axis corresponds to fractions with the top of the gradient to the left. A representative experiment of two performed is shown.

**Figure S15. Alteration in RPGs expression in T98G cells in response to serum starvation and DYRK1A depletion.**

Violin plots indicating the alterations in RNA expression levels of all human RPGs in T98G cells expressed as $log_2$FC in comparisons between cells serum- starved for 48 h and cells grown in normal conditions (Cycling). The data are shown for cells infected with a lentivirus expressing a shRNA Control (blue) and for cells infected with a lentivirus expressing a shRNA to DYRK1A (orange).

#### Figure S1

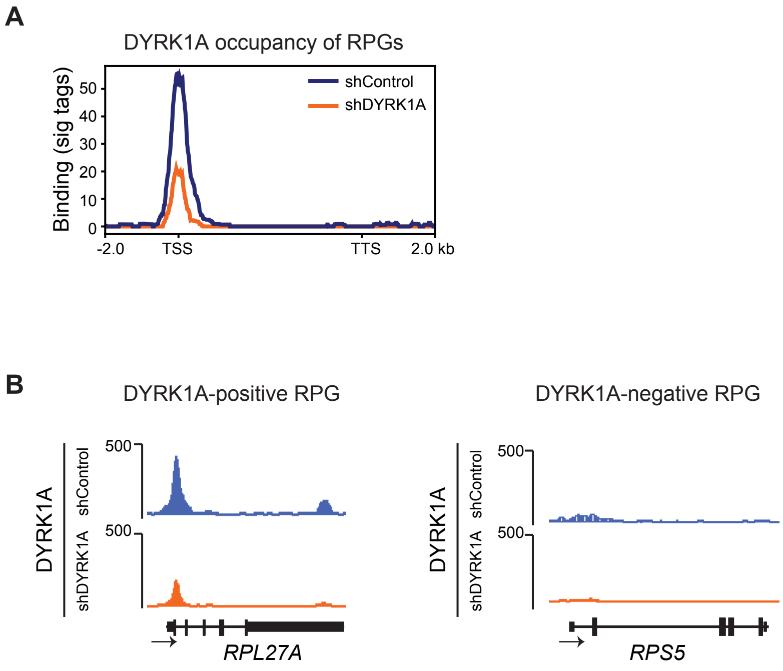
A
DYRK1A occupancy of RPGs
Binding (sig tags)
50
40
30
20
10
0
shControl
shDYRK1A
-2.0
TSS
TTS
2.0 kb
B
DYRK1A-positive RPG
DYRK1A-negative RPG
DYRK1A
shControl
shDYRK1A
500
RPL27A
RPS5

### Figure S2

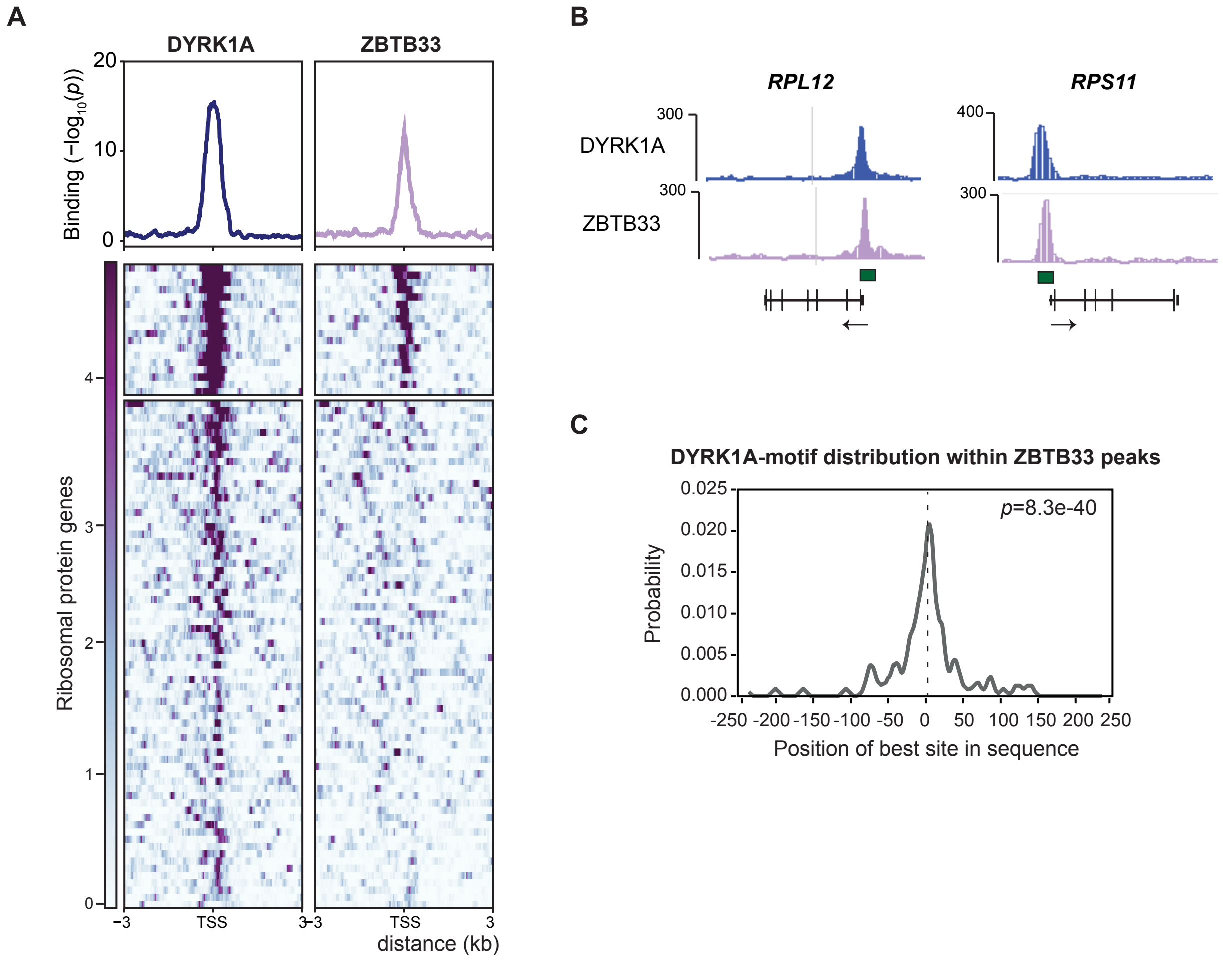

A
DYRK1A
ZBTB33
Binding (−log10(p))
20
10
0
Ribosomal protein genes
4
3
2
1
0
−3
TSS
3
-3
TSS
3
distance (kb)
B
RPL12
RPS11
DYRK1A
ZBTB33
300
300
400
300
C
DYRK1A-motif distribution within ZBTB33 peaks
p=8.3e-40
Probability
0.025
0.020
0.015
0.010
0.005
0.000
-250 -200 -150 -100 -50 0 50 100 150 200 250
Position of best site in sequence

#### Figure S3

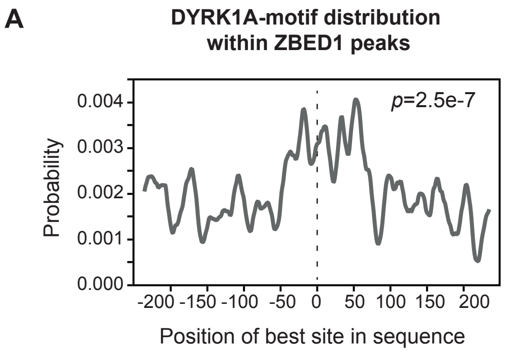
A
DYRK1A-motif distribution
within ZBED1 peaks
p=2.5e-7
Probability
0.004
0.003
0.002
0.001
0.000
-200 -150 -100 -50 0 50 100 150 200
Position of best site in sequence

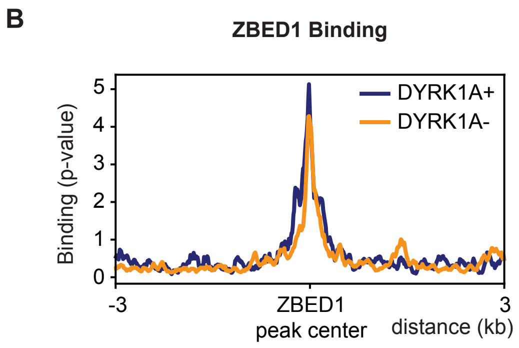
B
ZBED1 Binding
DYRK1A+
DYRK1A-
Binding (p-value)
5
4
3
2
1
0
-3
ZBED1
peak center
3
distance (kb)

Figure S4

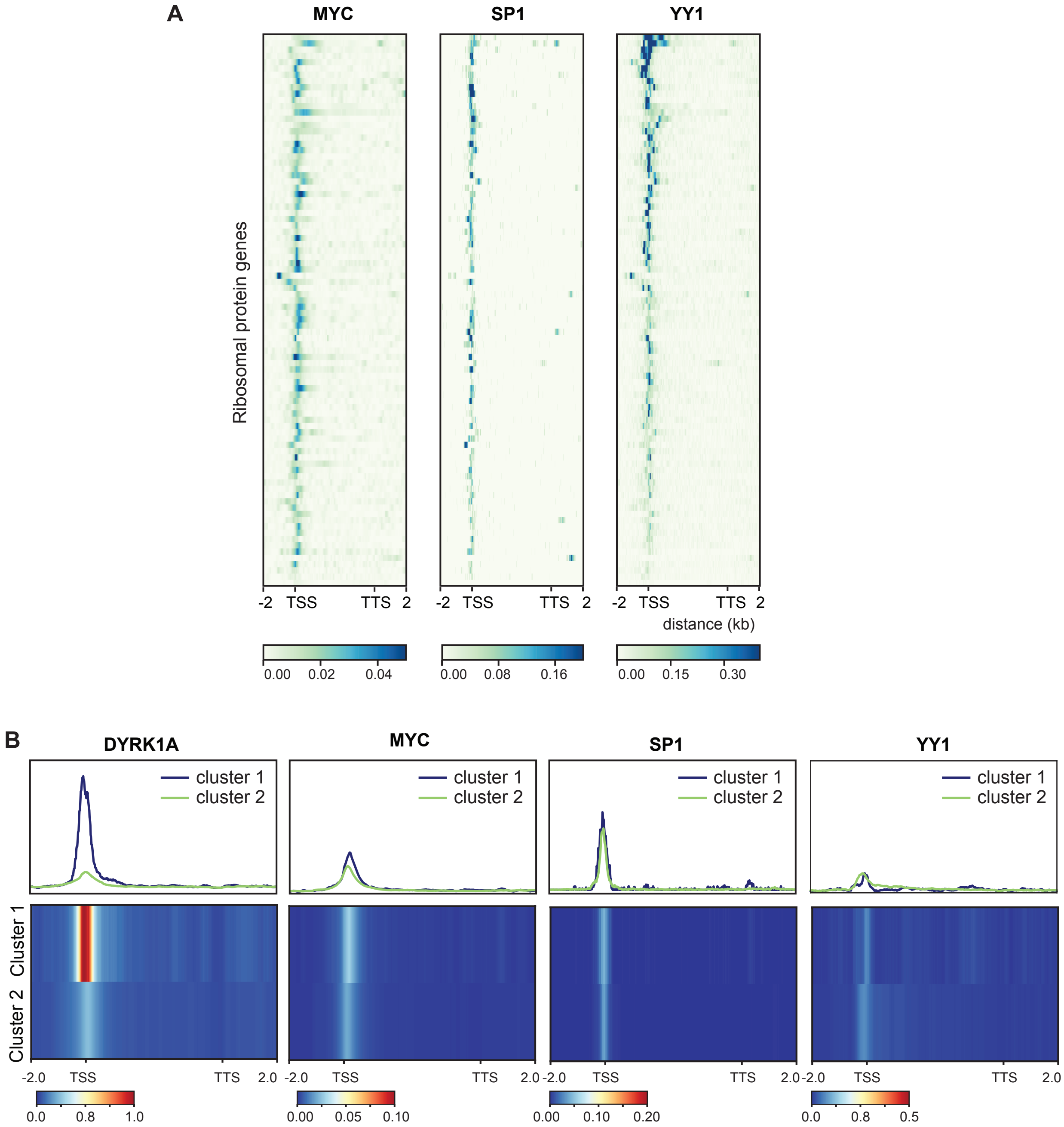
A
MYC
SP1
YY1
Ribosomal protein genes
-2 TSS TTS 2
distance (kb)
0.00 0.02 0.04
0.00 0.08 0.16
0.00 0.15 0.30
B
DYRK1A
MYC
SP1
YY1
cluster 1
cluster 2
Cluster 1
Cluster 2
-2.0 TSS TTS 2.0
0.0 0.8 1.0
0.00 0.05 0.10
0.00 0.10 0.20
0.0 0.8 0.5

### Figure S5

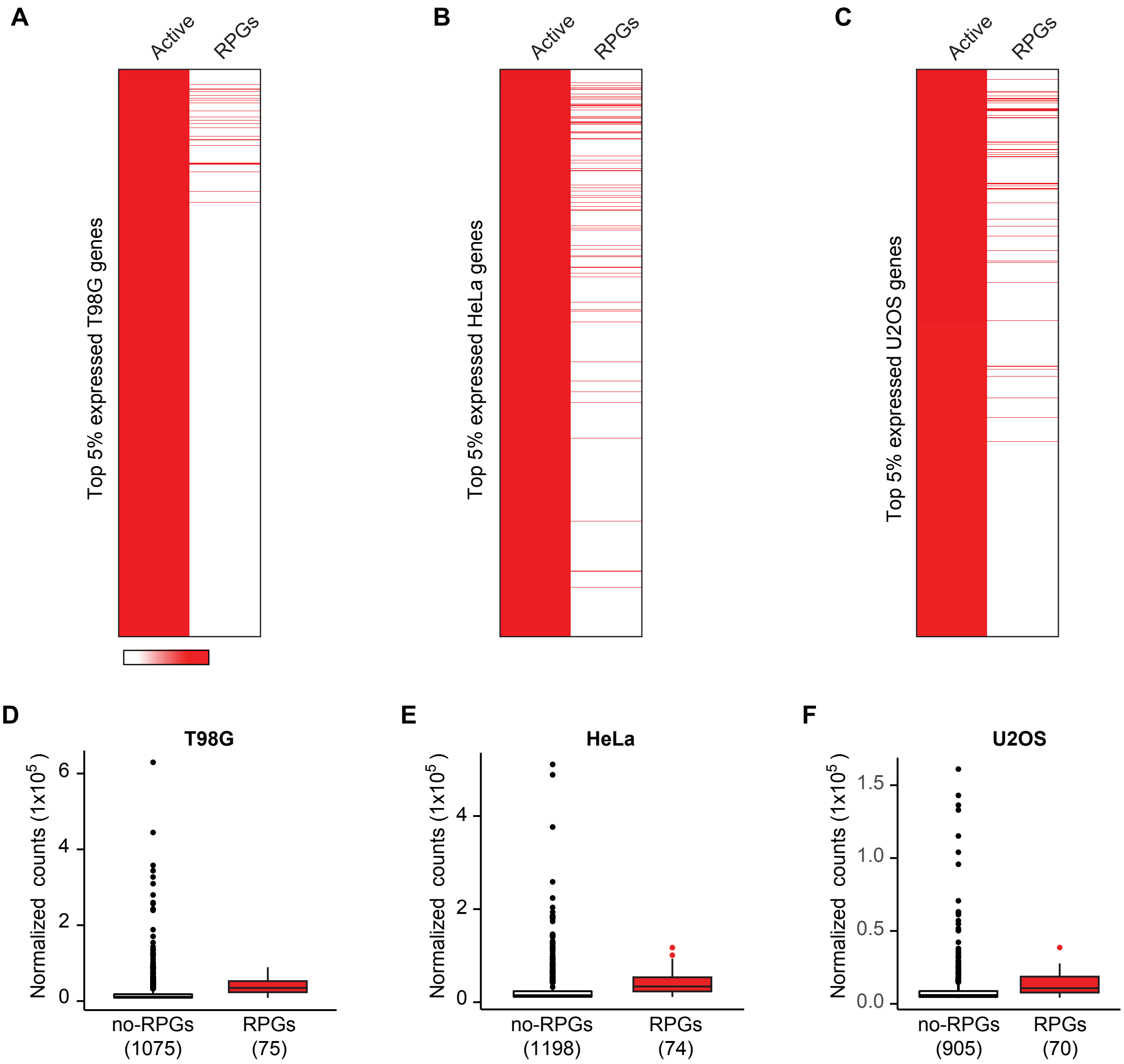
A
B
C
Active
RPGs
Top 5% expressed T98G genes
Top 5% expressed HeLa genes
Top 5% expressed U2OS genes
D
E
F
T98G
HeLa
U2OS
Normalized counts (1x10^5)
no-RPGs (1075)
RPGs (75)
no-RPGs (1198)
RPGs (74)
no-RPGs (905)
RPGs (70)
0
2
4
6
0.0
0.5
1.0
1.5

**Figure S6**

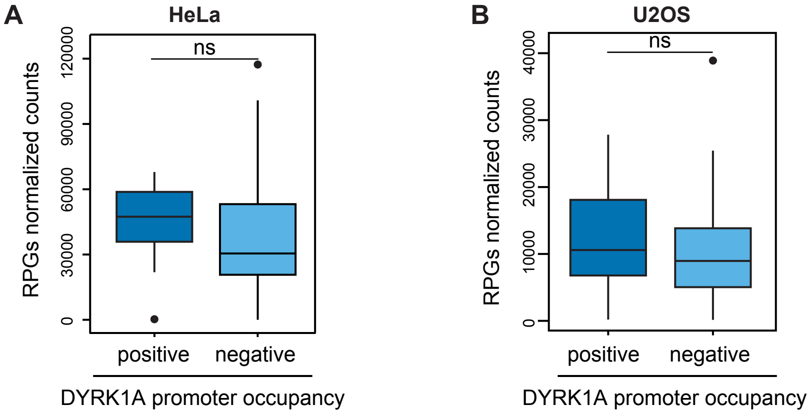
A
HeLa
B
U2OS
ns
RPGs normalized counts
0
30000
60000
90000
120000
10000
20000
30000
40000
positive
negative
DYRK1A promoter occupancy

### Figure S7

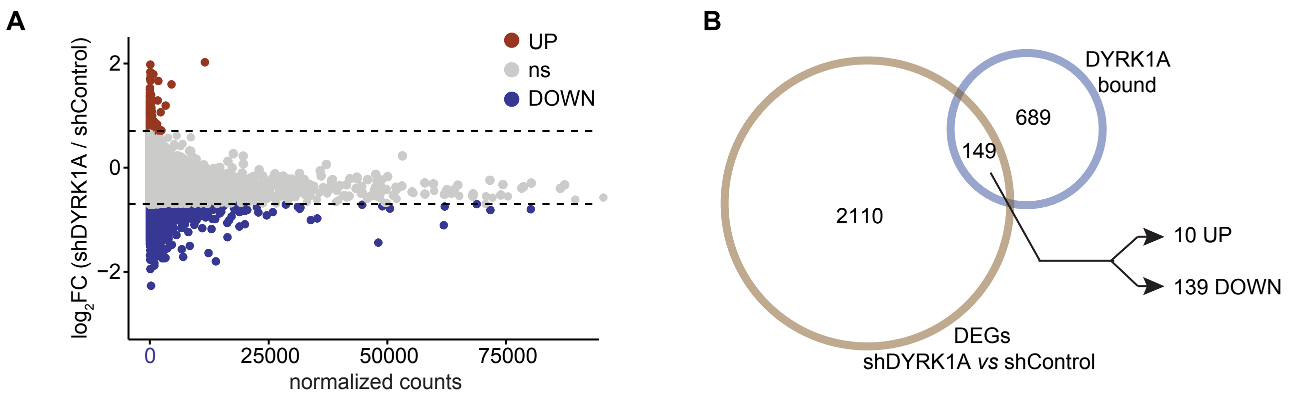
A
UP
ns
DOWN
log2FC (shDYRK1A / shControl)
2
0
−2
0
25000
50000
75000
normalized counts
B
DYRK1A
bound
689
149
2110
10 UP
139 DOWN
DEGs
shDYRK1A vs shControl

### Figure S8

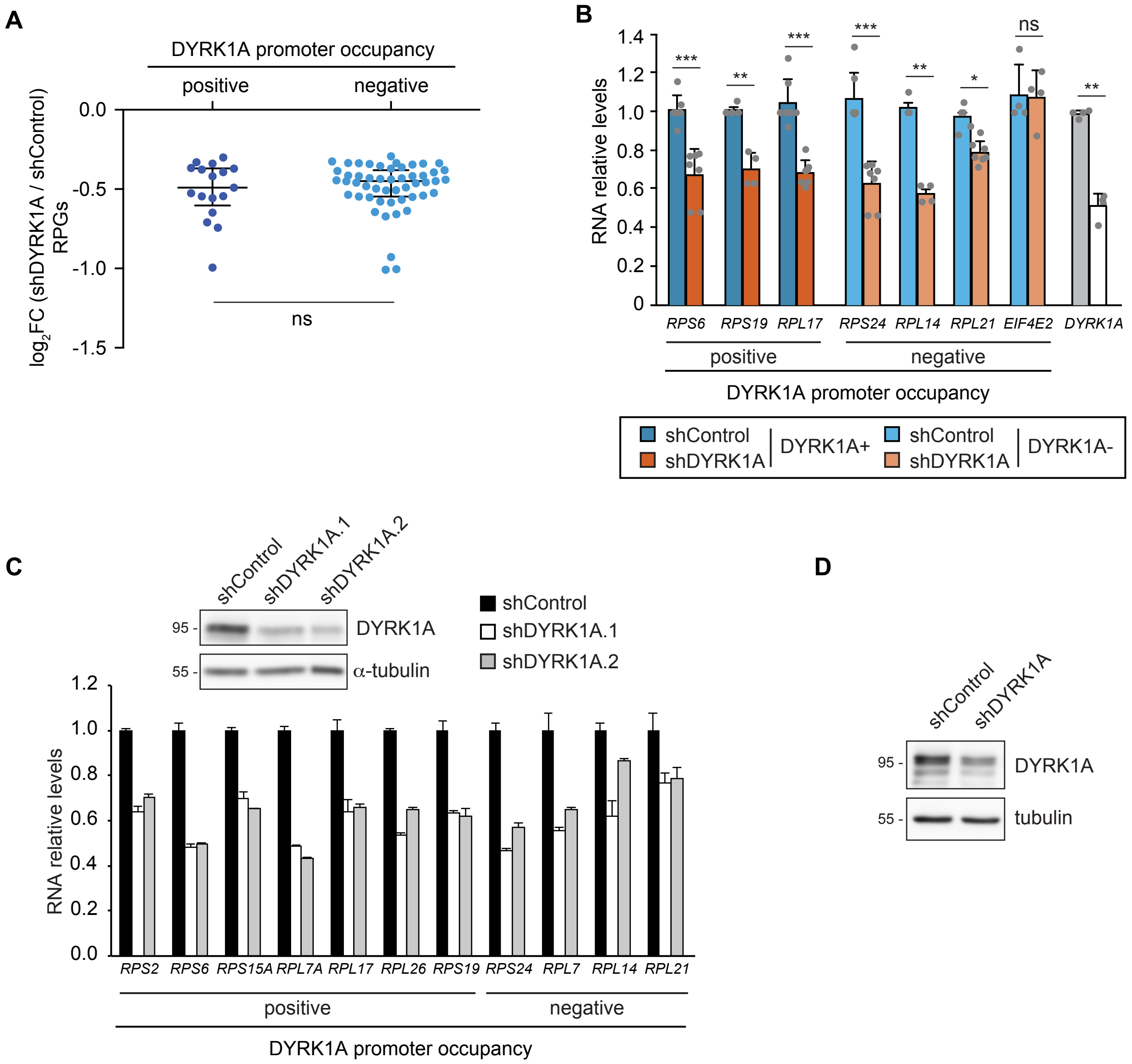

A
DYRK1A promoter occupancy
positive
negative
log2FC (shDYRK1A / shControl) RPGs
0.0
-0.5
-1.0
-1.5
ns
B
RNA relative levels
1.4
1.2
1.0
0.8
0.6
0.4
0.2
0
***
**
***
***
**
*
ns
**
RPS6
RPS19
RPL17
RPS24
RPL14
RPL21
EIF4E2
DYRK1A
positive
negative
DYRK1A promoter occupancy
shControl
shDYRK1A
DYRK1A+
shControl
shDYRK1A
DYRK1A-
C
shControl
shDYRK1A.1
shDYRK1A.2
95 -
DYRK1A
55 -
α-tubulin
shControl
shDYRK1A.1
shDYRK1A.2
RNA relative levels
1.2
1.0
0.8
0.6
0.4
0.2
0.0
RPS2
RPS6
RPS15A
RPL7A
RPL17
RPL26
RPS19
RPS24
RPL7
RPL14
RPL21
positive
negative
DYRK1A promoter occupancy
D
shControl
shDYRK1A
95 -
DYRK1A
55 -
tubulin

#### Figure S9

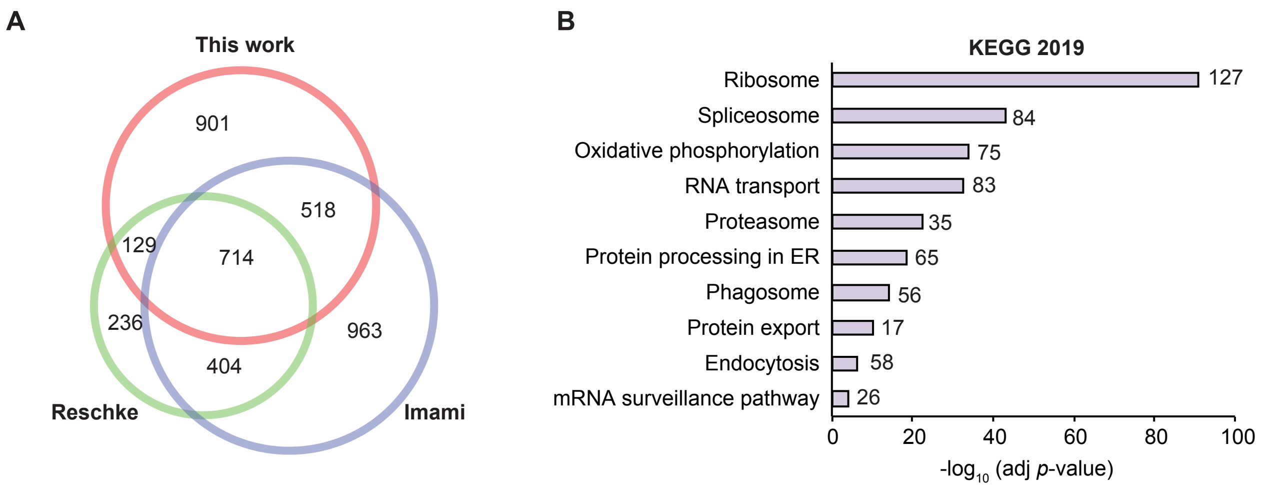
A
This work
901
518
129
714
236
963
404
Reschke
Imami
B
KEGG 2019
Ribosome
127
Spliceosome
84
Oxidative phosphorylation
75
RNA transport
83
Proteasome
35
Protein processing in ER
65
Phagosome
56
Protein export
17
Endocytosis
58
mRNA surveillance pathway
26
0
20
40
60
80
100
-log10 (adj p-value)

### Figure S10

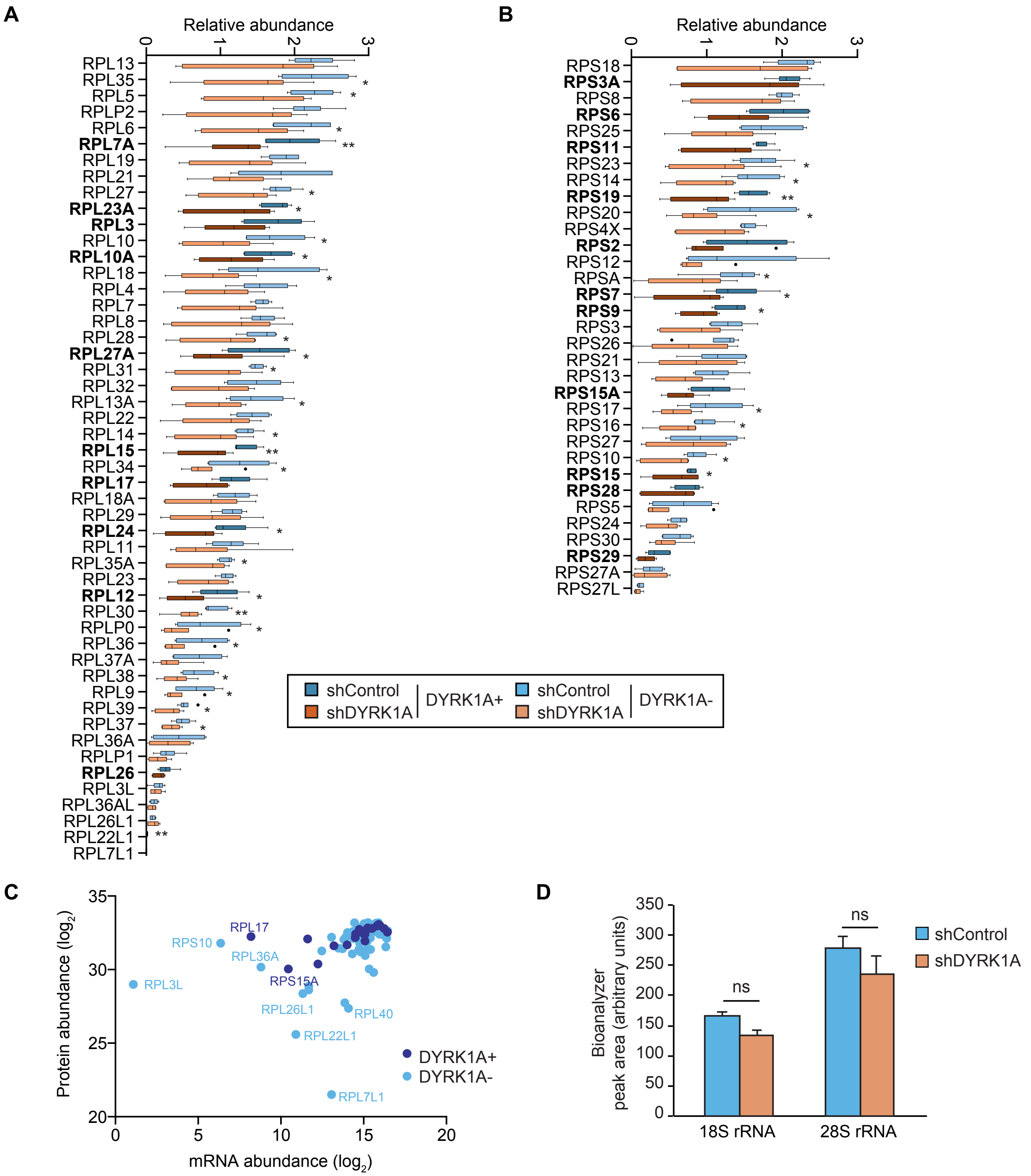
A
Relative abundance
0 1 2 3
RPL13
RPL35
RPL5
RPLP2
RPL6
RPL7A
RPL19
RPL21
RPL27
RPL23A
RPL3
RPL10
RPL10A
RPL18
RPL4
RPL7
RPL8
RPL28
RPL27A
RPL31
RPL32
RPL13A
RPL22
RPL14
RPL15
RPL34
RPL17
RPL18A
RPL29
RPL24
RPL11
RPL35A
RPL23
RPL12
RPL30
RPLP0
RPL36
RPL37A
RPL38
RPL9
RPL39
RPL37
RPL36A
RPLP1
RPL26
RPL3L
RPL36AL
RPL26L1
RPL22L1
RPL7L1
B
Relative abundance
0 1 2 3
RPS18
RPS3A
RPS8
RPS6
RPS25
RPS11
RPS23
RPS14
RPS19
RPS20
RPS4X
RPS2
RPS12
RPSA
RPS7
RPS9
RPS3
RPS26
RPS21
RPS13
RPS15A
RPS17
RPS16
RPS27
RPS10
RPS15
RPS28
RPS5
RPS24
RPS30
RPS29
RPS27A
RPS27L
shControl
shDYRK1A
DYRK1A+
shControl
shDYRK1A
DYRK1A-
C
Protein abundance (log2)
35
30
25
20
RPL17
RPS10
RPL36A
RPS15A
RPL3L
RPL26L1
RPL40
RPL22L1
RPL7L1
DYRK1A+
DYRK1A-
0 5 10 15 20
mRNA abundance (log2)
D
Bioanalyzer peak area (arbitrary units)
350
300
250
200
150
100
50
0
ns
ns
18S rRNA
28S rRNA
shControl
shDYRK1A

### Figure S11

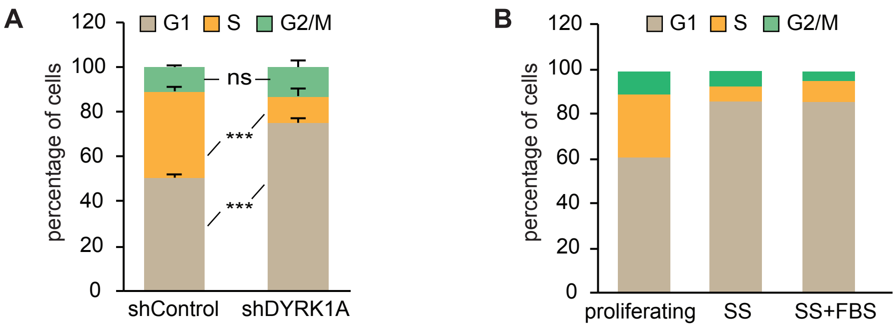
A
G1
S
G2/M
120
100
80
60
40
20
0
percentage of cells
ns
***
***
shControl
shDYRK1A
B
G1
S
G2/M
120
100
80
60
40
20
0
percentage of cells
proliferating
SS
SS+FBS

### Figure S12

A

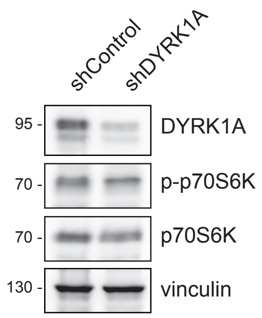

B

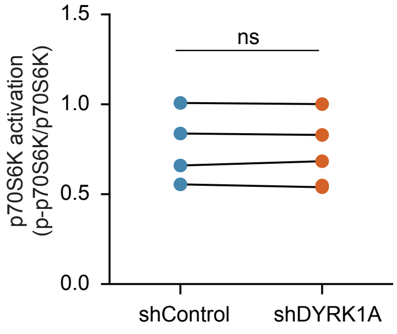

C

| | | shControl_1 | shControl_2 | shControl_3 | shDYRK1A_1 | shDYRK1A_2 | shDYRK1A_3 | tTest |
|---|---|---|---|---|---|---|---|---|
| Experiment 1 | RPS6 total | 6,34E+09 | 1,49E+10 | 4,40E+09 | 4,11E+09 | 3,50E+09 | 5,92E+09 | |
| | [R].LS**S**$^{236}$LRA**S**$^{240}$TSK.[S] | 6,23E+08 | 1,15E+09 | 3,94E+08 | 3,04E+08 | 3,09E+08 | 5,49E+08 | |
| | Ratio phosphpeptide/total RPS6 | 9,82E-02 | 7,74E-02 | 8,96E-02 | 7,40E-02 | 8,82E-02 | 9,28E-02 | 7,25E-01 |
| Experiment 2 | RPS6 total | 7,63E+09 | 2,25E+10 | 1,57E+10 | 1,13E+09 | 6,90E+08 | 1,26E+10 | |
| | [R].LS**S**$^{236}$LRA**S**$^{240}$TSK.[S] | 1,74E+07 | 1,43E+07 | 1,92E+07 | 1,52E+06 | 1,37E+06 | 1,67E+07 | |
| | Ratio phosphpeptide/total RPS6 | 2,29E-03 | 6,34E-04 | 1,22E-03 | 1,35E-03 | 1,99E-03 | 1,33E-03 | 9,37E-01 |

### Figure S13

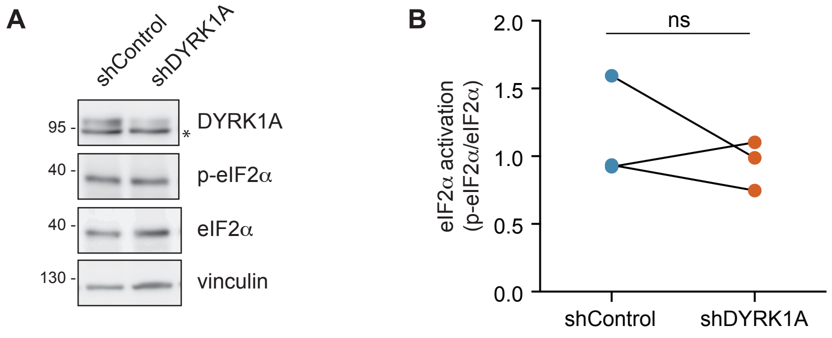
A
shControl
shDYRK1A
95 -
40 -
40 -
130 -
DYRK1A
*
p-eIF2α
eIF2α
vinculin
B
eIF2α activation
(p-eIF2α/eIF2α)
ns
2.0
1.5
1.0
0.5
0.0
shControl
shDYRK1A

**Figure S14**

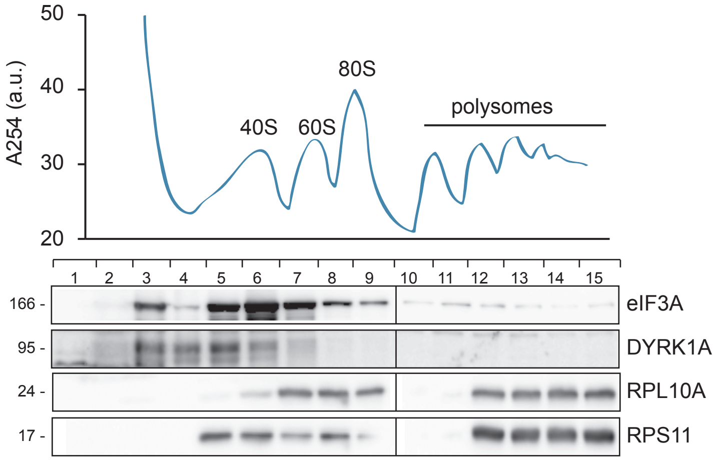
50
40
30
20
A254 (a.u.)
40S
60S
80S
polysomes
166 -
95 -
24 -
17 -
eIF3A
DYRK1A
RPL10A
RPS11

**Figure S15**

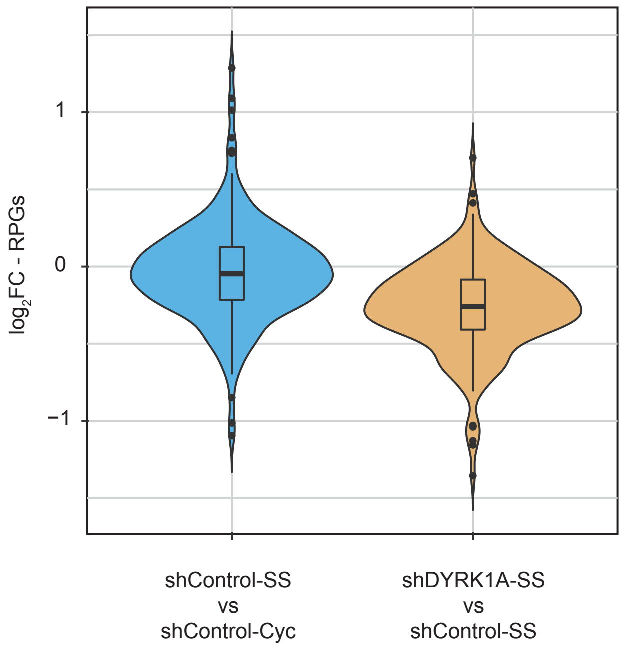

1
0
−1
$log_2FC$ - RPGs
shControl-SS
vs
shControl-Cyc
shDYRK1A-SS
vs
shControl-SS
